# Dynamics of mutators of arbitrary dominance in humans

**DOI:** 10.64898/2026.09.10.750760

**Authors:** Matin Saeidi, Guy Sella, Molly Przeworski, William Milligan

**Affiliations:** Department of Biological Sciences, Columbia University, New York, USA; Program for Mathematical Genomics, Columbia University, New York, USA; Irving Institute for Cancer Dynamics, Columbia University, New York, USA

## Abstract

Recent findings in humans and other species have revealed the presence of “mutator” alleles that increase germline mutation rate across the genome. Such mutators are expected to be selected against because of the additional deleterious alleles that they generate, to a degree that will depend on how much they increase the mutation rate in heterozygotes and homozygotes. To describe their dynamics, we develop a population genetic model of mutation rate modifiers with arbitrary dominance coefficients, in which fitness effects stem from additional germline mutations. We then use it to interpret findings for the seven human mutators identified to date. For six of the seven known mutators, the observed frequencies are well fit by the model and thus consistent with purifying selection arising solely due to their effects on germline mutation rates, although also consistent with a wide range of parameters; in two, the observed frequencies are readily explained by purely recessive fitness effects. The exception is a variant in *MUTYH*, which is more common than predicted under plausible parameters, for reasons that remain unclear. We also use the model to explore what types of mutators are most likely to be discovered in parents, based on identifying offspring with unexpectedly high numbers of *de novo* mutations. Although the two first mutators identified by this approach seem to be recessive, our modeling suggests that, for the same effect size, semi-dominant mutators are much more likely to be detected. These findings therefore imply that there are many more modifier sites with recessive effects than semi-dominant ones. More generally, our model provides a framework for interpreting properties of mutators in humans and other species and for learning about the genetic architecture of germline mutation rate variation.

## Introduction

Germline mutations are the ultimate cause of heritable disease and the substrate for evolution. Yet mutation rates themselves are evolving traits. For instance, mutation rates per year are lower in longer-lived species (reviewed in De Manuel et al. (2026)) and divergence in the mutation spectrum increases with phylogenetic distance in mammals (Goldberg & Harris 2022; Harris & Pritchard 2017; Lee et al. 2025; Beichman et al. 2023; Talenti et al. 2025). Within humans, variation in the number of *de-novo* mutations (DNMs) is heritable (Sasani et al. 2019; Kaplanis et al. 2022). These observations indicate that there must be genetic variants that modify germline mutation rates in humans.

Such modifier variants will evolve subject to natural selection and genetic drift. A variant that increases germline mutation rates, i.e., a mutator, becomes transiently linked to the DNMs that it causes. As DNMs are deleterious on average (Eyre-Walker & Keightley 2007), mutators are expected to be under purifying selection (Kimura 1967; Kondrashov 1995; Lynch 2008). The fate of a mutator depends on the strength of selection acting against it. If selection is strong relative to the effects of drift, a mutator will be short-lived and kept at low frequencies. Conversely, a mutator under weak selection may ascend to high frequencies and even reach fixation in the population.

Differences in the strength of purifying selection against mutators may lead to differences in mean mutation rates across species. All else being equal, the “drift-barrier” hypothesis predicts that sexually-reproducing species with larger effective population sizes will more effectively purge mutators and thus evolve lower mutation rates per generation (Lynch 2010; Lynch 2011; Sung et al. 2012). This prediction is supported by comparisons among distantly related taxa, across which effective population sizes vary by many orders of magnitude (Sung et al. 2012; Lynch et al. 2023; Wang & Obbard 2023; Bergeron et al. 2023). However, the drift-barrier hypothesis assumes that there exist mutators with effect sizes that transition from effectively neutral to strongly selected as effective population size increases. This assumption may not hold among closely related taxa (e.g., primates or even mammals), between which there is little variation in effective population size (Prado-Martinez et al. 2013; Bergeron et al. 2023). Moreover, changes in population size are often accompanied by changes in life-history traits, notably generation time (Chao & Carr 1993), that can weaken or even invert the relationship between population size and mutation rate (De Manuel et al. 2026; Zhu et al. 2025). As such, the extent to which variation in mean mutation rates among closely related taxa reflects differences in the efficacy of purifying selection against mutators is unclear (Weinstein & Roy 2026).

To date, most theory focuses on modeling differences in mutation rates across species, and does not consider their dynamics within a species (Kimura 1967; Lynch 2010; Zhu et al. 2025). Mutators that are strongly selected in small populations may never fix and thus are unlikely to contribute to variation between species (Lynch 2011). However, such mutators may transiently increase mutation rates during their sojourn within a population (Milligan et al. 2022). Theoretical models of how mutation rates evolve within a population have received little attention: Lynch (2008) examined the equilibrium frequency of mutators but ignored the effects of drift in shaping variation within a population, while Milligan et al. (2022) examined the distribution of mutator frequencies under mutation-selection-drift balance but assumed mutators act additively.

The evolution of mutation rates is further complicated by potential selection against the pleiotropic effects of mutators. For example, if mutators increase the rate of somatic mutations and thus the risk of cancer, they may be under stronger purifying selection than their germline effects alone would predict (Lynch 2008). Conversely, selection against mutators could also be weaker than expected, if mutation rates are under (indirect) stabilizing selection due to trade-offs between lowering the mutation rate and organismal constraints such as developmental timing (Kimura 1967; Kondrashov 1995; Dawson 1998). The role of pleiotropy and constraint in shaping the evolution of germline mutation rates remains poorly understood (Zhu et al. 2025; De Manuel et al. 2026).

Recently, however, a number of naturally-occurring mutators have been identified in humans and other mammals, which may allow us to learn more about the evolution of mutators and germline mutation rates (for earlier work in microorganisms, see, e.g., Jyssum (1960), Gross and Siegel (1981), Gou et al. (2019), and Jiang et al. (2021)). Notably, a pedigree-based study of DNMs in over ten thousand human trios led to the discovery of twelve offspring carrying many more *de novo* point mutations than expected (henceforth referred to as a “proband”) (Kaplanis et al. 2022). In two such cases, the fathers were homozygous for a disruptive variant (loss of function or deleterious missense mutation) in *XPC* and *MPG*, known DNA repair genes. Five additional mutators have been identified by sequencing trios ascertained on a parent carrying a missense variant in either one of two DNA polymerase genes (*POLE* and *POLD1*) or the base-excision repair gene, *MUTYH* (Sherwood et al. 2023; Young et al. 2024). Finally, disruptions of *REV1* and *LIG1* were found to be associated with increased point mutation rates in humans using a burden test, though the causal variant(s) remain unknown (Getseva et al. 2026). Beyond humans, a mutator in *MBD4* was identified in a rhesus macaque dam, whose offspring carried abnormally high numbers of germline point mutations (Stendahl et al. 2023). In mice, quantitative trait locus mapping of variation in C *>* A mutation rates revealed a cluster of five alleles in *MUTYH* that decrease the rate of such mutations (relative to the ancestral alleles) (Sasani et al. 2022; Sasani et al. 2024).

By contrast, mapping efforts in humans using association studies have failed to identify common variants associated with germline mutation rates (Kaplanis et al. 2022; Garcia-Salinas et al. 2025). The lack of signal suggests that mutators with detectable effects are not identified because they are rare, owing to strong purifying selection. Nonetheless, the magnitude of mutator effects on germline mutation rates varies markedly among discovered examples. For instance, the offspring of a father homozygous for the *XPC* mutator inherited ~6.5 fold more DNMs than expected given the ages of their parents (Kaplanis et al. 2022), whereas offspring of parents that carried a missense mutation in *POLE* only inherited ~2.3 fold more DNMs than controls (Sherwood et al. 2023). In turn, missense and LoF mutations in *REV1* increased the overall mutation rate by only 16% on average and, similarly, disrupting mutations in *LIG1* increased the CpG *>* TpG mutation rate by 24% on average (Getseva et al. 2026). One caveat to the interpretation of effect sizes at *REV1* and *LIG1* is that burden tests consider sets of variants in aggregate, so a subset of the variants could have larger effects.

Intriguingly, these examples also vary in their dominance effects. At *POLE* and *POLD1*, mutation rates are significantly increased in heterozygous carriers (Sherwood et al. 2023); the same seems to be true for *REV1* and *LIG1* (Getseva et al. 2026). In contrast, the mutators in *MUTYH* appear to act recessively: mutation rates are significantly increased in compound heterozygotes but not in simple heterozygotes (Young et al. 2024). Consistent with this mode of action, *MUTYH* is one of the few disease genes classified as recessive by clinicians in which loss-of-function (LoF) allele frequencies are compatible with a model of strictly recessive fitness effects and in which a single copy of a LoF allele has no detectable effects on the quantitative traits examined (Judd et al. 2026). The dominance of the human mutators in *XPC* and *MPG* is unknown, having only been reported for one homozygous parent each, but presumed to be recessive (Kaplanis et al. 2022). In line with this assumption, only humans with two disrupted copies of *XPC* develop xeroderma pigmentosum (Lehmann et al. 2011), and *XPC* LoF alleles act as recessive mutators in mice (Miccoli et al. 2007).

Motivated by these observations, we generalize the mutator model developed in Milligan et al. (2022) to allow for arbitrary dominance coefficients rather than assume semi-dominance. We rely on this model to ask whether the estimates of frequencies, effects on germline mutation rates, and dominance of known mutators are consistent with selection acting only through their germline effects. We also ask about the type of mutators that we should expect to find using current pedigree-based approaches, in terms of effect sizes and dominance coefficients.

### The Model

We assume that an individual’s genotype consists of two parts: *G* biallelic sites, a fraction *f* of which are under strong selection, and *M* biallelic modifier sites that determine the mutation rate (see Table 1 for a summary of notation). The remaining (1 *− f*)*G* sites are neutral and have no effect on fitness. We further assume fitness combines multiplicatively over the selected sites and that each deleterious allele reduces fitness by a factor of *s*_*het*_, where *s*_*het*_ *>* 0. Thus, an individual carrying *i* deleterious alleles has absolute fitness (1 *− s*_*het*_)^*i*^. Our results should hold even when fitness effects vary across sites (see Milligan et al. (2022); Figure S4), in which case *s*_*het*_ can be interpreted as the average fitness effect at selected sites.

**Table 1:** Parameters and the values considered.

| Parameter | Meaning | Value |
| --- | --- | --- |
| $N_e$ | Effective population size | $2 \cdot 10^4$ or varies |
| $\mu$ | Mutation rate at modifier sites | $1.25 \cdot 10^{-8}$ /bp/gen |
| $\hat{u}$ | Estimated point mutation rate | $1.25 \cdot 10^{-8}$ /bp/gen |
| $u_0$ | Baseline mutation rate at selected sites | $\hat{u} - M\mathbb{E}[\Delta u]$ |
| $G$ | Haploid genome size in basepairs | $3 \cdot 10^9$ |
| $f$ | Fraction of genome under selection | $8 \cdot 10^{-2}$ |
| $s_{het}$ | Heterozygous selection coefficient of a deleterious allele | $5 \times 10^{-4}$ |
| $M$ | Number of modifier sites | varies |
| $\phi$ | Increase in mutation rate in mutator homozygotes relative to anti-mutator homozygotes | varies |

Our mutation model generalizes the model from Milligan et al. (2022) to allow for any dominance coefficient at modifier sites. We first assume a minimum mutation rate of *u*_0_ per site per generation, which is the rate obtained when DNA repair and replication are optimized. We further assume modifier sites switch between two states: anti-mutator and mutator. A mutator increases the mutation rate across the genome by *ϕ* in homozygotes and by *hϕ* in heterozygotes relative to those homozygous for the anti-mutator. For now, we consider that mutators at separate modifier sites act independently; that is, we do not allow for epistasis or compound heterozygosity. Thus, the mutation rate, *u*, of a person is

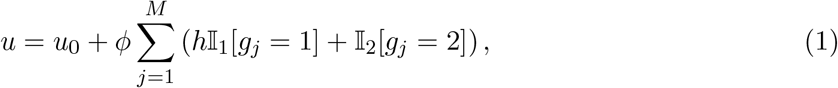

where *g*_*j*_ ∈*{*0, 1, 2*}* is the number of mutators that the person carries at site *j* and I _1_ and I _2_ are indicator functions that describe whether the person is heterozygous or homozygous for the mutator at a given site. Our model ignores known age and sex effects in vertebrates (Ségurel et al. 2014; De Manuel et al. 2022; Bergeron et al. 2023), because the evolutionary dynamics at modifier sites are well approximated in terms of the sex- and age-averaged rates. Our main results should hold under more complex mutation models, e.g., with a distribution of mutator effect sizes (see Milligan et al. (2022); Figure S5) and dominance coefficients or when mutators affect only a subset of the genome.

We first model the evolution of mutation rates in a diploid, panmictic population of constant size *N*_*e*_, with non-overlapping generations. In each generation, individuals are randomly chosen as parents with probabilities proportional to their fitness (i.e., according to Wright-Fisher sampling with fertility selection). These individuals then produce gametes with free recombination among selected and modifier sites, Mendelian segregation, and mutations at both the modifier and selected sites. We assume that the mutational input per generation per selected site is sufficiently low (2*N*_*e*_*u≫*1) and that deleterious alleles are sufficiently rare (2*N*_*e*_*s*_*het*_ *≪* 1) such that we can apply the infinite sites approximation and ignore reverse mutations. The number of deleterious alleles added per gamete per generation is then Poisson distributed with mean *Gfu*. We further assume that the mutation rate at modifier sites per generation, *µ*, is symmetric and constant, i.e., it is not affected by mutators (see Milligan et al. (2022) for the potential effects of violating this assumption). The resulting gametes combine randomly to produce the next generation.

In what follows, we parameterize our model using reasonable values for humans (Table 1). When modeling a population of constant size, we use *N*_*e*_ = 20,000, the ancestral effective population size estimated by Schiffels and Durbin (2014). Later, we allow the population size to vary according to inferred South Asian (SAS), African/African American (AFR) or non-Finnish European (NFE) demographic histories (see Supplementary Section S3.5 for details). For the mutation parameters, we allow the number of modifier sites, *M*, and the effect size of mutators, *ϕ*, to vary. In turn, we set *u*_0_ such that the expected mutation rate conditional on *ϕ* and *M* matches the estimated mutation rate per base per generation in humans, *û* = 1.25 *·* 10^*−*8^ (Jónsson et al. 2017). When we apply our model to known mutators, we use estimates of *µ* taken from the Roulette model (Seplyarskiy et al. 2023) and assume forward and backward mutation rates are the same; otherwise, we assume that *µ* = *û*. For the selection coefficients, we assume a haploid genome length of *G* = 3 *×* 10^9^ base pairs (Nurk et al. 2022) and rely on estimates that roughly *f* = 8% of the human genome is under strong selection (Davydov et al. 2010; Ponting & Hardison 2011; Ward & Kellis 2012; Rands et al. 2014). We further assume that, in this subset of the genome, mutations have a fixed fitness effect of *s*_*het*_ *≈* 5 *×* 10^*−*4^ (Agarwal & Przeworski 2021). We explore alternative values of *s*_*het*_ in Figures S2 and S7.

### The dynamics of mutators at steady state

We use our model to study the dynamics of mutators at steady state. Specifically, we investigate how the dominance and effect size of mutators impact their expected frequencies and their contribution to the mean mutation rate (averaged across the population). To this end, we first approximate the expected selection coefficient of a mutator. Selection against a mutator arises from its linkage with the additional deleterious alleles that it generates. We approximate the expected additional number of mutations that become linked to a mutator (with effect size *ϕ*, dominance coefficient *h* and frequency *q*) each generation as

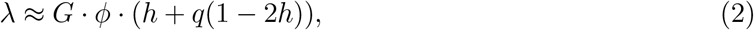

where the 1 *−* 2*h* term reflects the deviation from additivity (Kimura 1960). Assuming free recombination, such mutations will remain linked to the mutator for an average of two generations (Kimura 1967). A proportion *f* of these mutations are strongly selected against, so the selection coefficient of the mutator is approximately

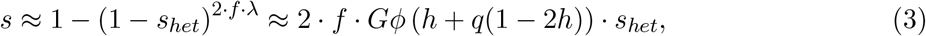

where the latter approximation assumes that 2*fε · s*_*het*_ *≪* 1. All of the above approximations assume that the number of additional deleterious alleles linked to the mutator nears its asymptotic value, 2*ε*, much faster than the frequency of the mutator changes (Figure S1); for further details, see Supplementary Section S2.2.

We can now approximate the stationary frequency distribution of a mutator, based on the first two moments of change in mutator frequency per generation (Ewens 2004, Chapter 4). The expected change in mutator frequency is well approximated by

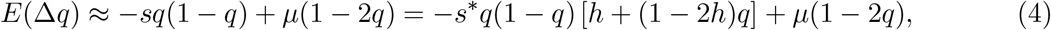

where the first term is the change due to selection with

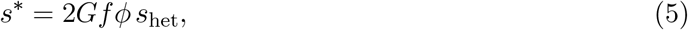

and the second term is the change due to mutation. The change due to selection takes the additive form because, on average, each copy of the mutator is linked to the same excess number of deleterious alleles. However, equation 4 exactly matches the standard expression for a deleterious allele with a constant selection coefficient, *s*\*, and dominance coefficient, *h* (Kimura (1960), see Supplementary section S3.2). The second moment of change in allele frequency follows the standard drift term,

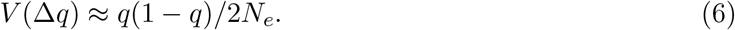

Given these moments, the stationary distribution takes the form

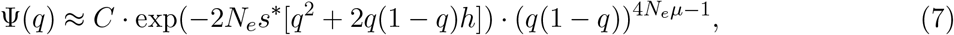

where *C* is a normalizing constant such that 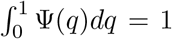 (see Supplementary section S2.3 for details).

We calculate the expected frequency of a mutator and the corresponding increase in the population mean mutation rate by integrating over the stationary distribution. The first two moments of mutator frequency are

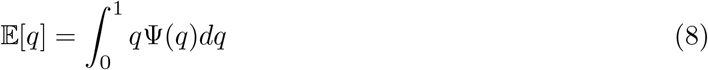

and,

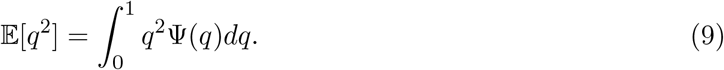

Assuming Hardy-Weinberg equilibrium, the expected increase in the population mean mutation rate due to a mutator at frequency *q* is

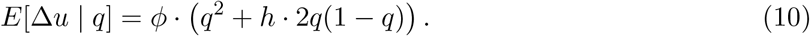

The expected increase in the population mean mutation rate owing to this mutator, without conditioning on a given frequency, is therefore

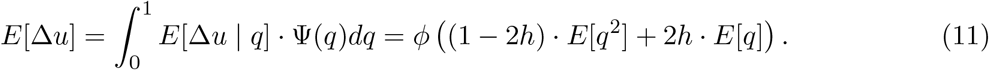

In our model, in which all mutators have the same dominance and effect size and evolve independently, the increase in the mean mutation rate from *M* modifier sites is simply

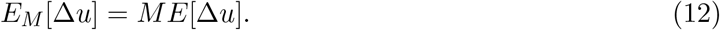

Similar results would be obtained if mutators varied in their effect sizes or dominance as we could also integrate Eq. 11 over distributions of *ϕ* and *h*. While assuming independence is sensible for mutators in distinct genes, it may not be for those that arise within the same gene. For example, compound heterozygotes may phenocopy individuals homozygous for a mutator, which could expose rare recessive mutators to selection (Clark 1998). To consider this case, we use simulations to estimate the frequency distribution.

The dynamics of mutators at steady state are similar to that of other deleterious alleles at mutation-selection-drift balance (Figure 1). In particular, they exhibit three regimes based on the scaled selection parameter, 2*N*_*e*_*s*\*: (i) The effectively neutral regime in which the selection against a mutator is much weaker than genetic drift (2*N*_*e*_*s*\* *≪* 1). In this regime, the expected frequency of the mutator is approximately 1/2, as it is equally likely to be at fixation or loss. The increase in the mean mutation rate then scales linearly with effect size, *ϕ*. (ii) The nearly neutral regime in which the effects of selection and genetic drift are comparable (2*N*_*e*_*s*\* *≈* 1). In this regime, a mutator may reach fixation but does so with lower probability than in the effectively neutral regime. The impact of modifiers on the mean mutation rate is maximized in this range (akin to the maximization of genetic load from deleterious alleles in this range (Kimura et al. 1963; Agrawal & Whitlock 2012)). Nonetheless, a single modifier site is expected to increase the mean mutation rate by less than 1% even if *s*_*het*_ is well below our assumed value (Figure S2). (iii) The strongly selected regime, in which selection is much stronger than drift (2*N*_*e*_*s*\* *≫* 1). In this regime, mutators almost never reach fixation. When partially dominant, the expected frequency of a mutator is inversely proportional to effect size; when recessive, the same applies to the expected frequency squared. In both cases, there is an approximately constant increase in the mean mutation rate throughout a broad range of effect sizes. At very large effect sizes, however, the linear relationship between mutator effect size and fitness breaks down, resulting in a slight increase in the mean mutation rate (Supplementary Section S2.2).

**Figure 1:**
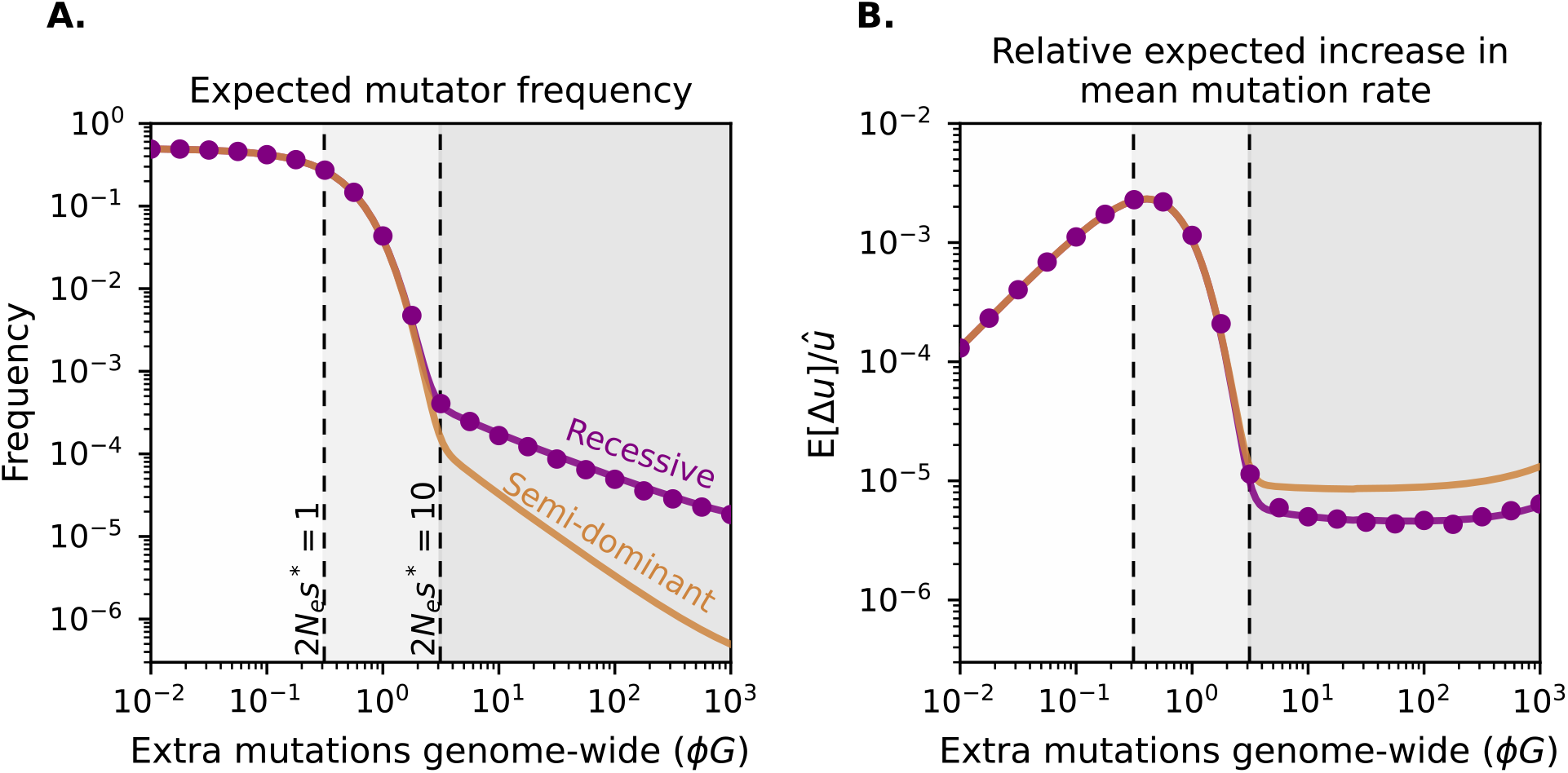
Expected mutator frequency (**A**) and contribution to the mean mutation rate (**B**) as a function of mutator effect size. We report effect sizes in units corresponding to the expected additional mutations genome-wide per generation, *ϕG* and the increase in mutation rate relative to the estimated human point mutation rate, i.e., *E*[Δ*u*]*/û*. Solid lines correspond to the analytical expectations calculated from Eq. 8 (in A) and Eq. 11 (in B). Circles correspond to the simulated mean estimated from 50 replicates; error bars indicating *±*2 standard errors are too small to be visible and thus not shown. All results assume *N*_*e*_ = 2,000, with other parameters scaled such that the compound values 2*N*_*e*_*s*_*het*_, 2*N*_*e*_*µ*, and 2*f Gs*_*het*_ retain plausible values in humans (i.e., those corresponding to *N*_*e*_ = 20,000 in Table 1). Dashed black vertical lines mark boundaries between selection regimes defined by the scaled selection parameter: effectively neutral (2*N*_*e*_*s*\* *<* 1), nearly neutral (1 *<* 2*N*_*e*_*s*\* *<* 10), and strongly selected (2*N*_*e*_*s*\* *≥* 10).

Only when selection against mutators is strong (i.e., 2*N*_*e*_*s*\* *≫* 1) does dominance affect the frequency of mutators and thus their contribution to the mean mutation rate. In Figure 1, we compare semi-dominant and recessive mutators with the same effect in homozygotes (i.e., *ϕG*). In the strongly selected regime, the differences between these cases arise for the same reasons that they arise for deleterious mutations in standard population genetic models (Haldane 1937; Simons et al. 2014; Balick et al. 2015; Henn et al. 2015). Because semi-dominant mutators have effects in heterozygotes, purifying selection acts on them regardless of frequency, and all segregating copies contribute to the increase in mean mutation rate. In contrast, when rare, recessive mutators do not experience effective selection and do not affect the mean mutation rate. As a result, in the strongly selected regime, recessive mutators segregate at higher frequencies than semi-dominant ones but cause a smaller increase in the mean mutation rate on average. In the weakly selected regime, by contrast, the effect of mutators on the mean mutation rate is instead dominated by the expected time that they spend at fixation, on which dominance has little effect.

### Does this model readily predict observed mutator frequencies?

Next, we ask whether our model readily gives rise to the observed frequencies of known mutator alleles, assuming a realistic demographic model and other nuisance parameters. To do so requires an estimate of the effect size of the mutator. For each, we therefore estimate the average-fold increase in the number of DNMs passed on by carriers relative to an expectation that accounts for parental sex and age effects on germline mutation rates (Jónsson et al. 2017). We then use this fold increase to calculate the expected number of DNMs that would be passed on by a carrier, averaged across parental ages and sex (see Supplementary Section S4.1). We assume that the strength of selection that has typically acted against the mutator reflects this average effect size.

Our effect size estimates suggest that mutators discovered to date are likely to have evolved under strong selection. Specifically, based on the ancestral effective population size for humans (*N*_*e*_ = 20, 000) and our assumed mean fitness effect of a new mutation (*fs*_*het*_ = 4 *·* 10^*−*5^), mutators that cause homozygous parents to pass on more than ~3 additional DNMs on average should experience strong selection (2*N*_*e*_*s*\* *>* 10). The mutators in *XPC, MPG, MUTYH, POLE*, and *POLD1* have effect sizes much greater than this value and thus would be strongly selected against even if *fs*_*het*_ is an order of magnitude smaller than we assume. In contrast, disrupting variants in *REV1* and *LIG1* have average effect sizes that place them at the boundary of the strong selection regime, such that mutators in these genes may only be weakly selected against if *fs*_*het*_ is much smaller than we assume.

In evaluating whether our model predicts the observed frequencies of mutators, we focus on cases mapped to single alleles, and thus exclude the disrupting variation in *REV1* and *LIG1* identified by a burden test (Getseva et al. 2026). For each mutator, we simulate its predicted frequency distribution (see supplementary sections S3.3 and S3.4 for a description of simulations). We vary the selection coefficient, *s*\*, between *ŝ*\**/*50 and 50*ŝ*\*, where *ŝ*\* is our estimate based on the estimated effect size of the mutator (reported in Tables 2 and S3) and assuming *fs*_*het*_ = 4 *·* 10^*−*5^. The remaining parameters are held constant. For each mutator, we use the mutation rate for the corresponding mutation type estimated in Seplyarskiy et al. (2023). We assume equal forward and backward mutation rates, but this assumption has little effect on our results (Figure S9). We further assume mutators in *POLD1* and *POLE* are semi-dominant and those in *MUTYH, XPC*, and *MPG* are recessive. For two genes, we allow the mutator phenotype to arise, and thus selection to act, in compound heterozygotes: for *XPC*, we assume any LoF mutation, including frame-shifting indels, can cause the mutator phenotype, whereas for *MUTYH*, we assume that only individuals homozygous or compound heterozygous for the three known missense variants express the mutator phenotype. Allowing for more or less compound heterozygosity has little effect on our results (Figure S5).

**Table 2:** Known human mutators and their estimated effects on the germline mutation rate of the parent with the mutator.

| Gene | Mutator | Effect on germline mutation rate |  |  |
| --- | --- | --- | --- | --- |
|  | Variant frequency | Dominance | Fold increase in offspring | Extra DNMs genome-wide |
| *XPC* | $1.61 \times 10^{-5}$ | Unknown | 7.65 | 234 |
| *MPG* | $1.75 \times 10^{-4}$ | Unknown | 4.62 | 127 |
| *POLE* | $6.20 \times 10^{-7}$ | Dominant | 4.34 | 117 |
| *POLD1* | 0 in 1,599,574 chromosomes | Dominant | 2.64 | 58 |
| *MUTYH* | $^a 1.92 \times 10^{-3}$<br>$^b 4.73 \times 10^{-3}$<br>$^c 2.50 \times 10^{-4}$ | Recessive | 1.71 | 25 |

Because the mutators were discovered by surveying people of recent European genetic ancestry, which may bias the discovery towards variants at higher frequencies in that group, we use allele frequencies from the largest non-European population designation in gnomAD v4.1.1, namely people of recent South Asian ancestry (Guez et al. 2026). In doing so, we assume that mutators found in both European and South Asian groups are not identical-by-descent but rather arose from independent (i.e., recurrent) mutations, which is likely given their low frequencies (1000 Genomes Project Consortium 2015; Biddanda et al. 2020; Stolyarova et al. 2025). Thus, the frequency estimate in the South Asian group can be considered an independent estimate to that in Europeans. To model their demographic history, we rely on estimates from Kar et al. (2026).

For six out of the seven mutators tested, the predicted frequency distribution includes the observed value, at *ŝ*\* (Figure 2A). However, the likelihood of the data is flat throughout the entire range of selection coefficients considered (Figure 2B–F, Supplementary Section S5), and we cannot reject neutrality for any of the mutators (Table S6). There is little information carried by one mutator per gene (three for *MUTYH*), in part because these mutators are young. Simulations suggest that the median age of a neutral, derived allele is ~26 generations, which provides insufficient time for selection to generate frequency differences between neutral and deleterious alleles unless selection is nearly lethal (Figure S10).

**Figure 2:**
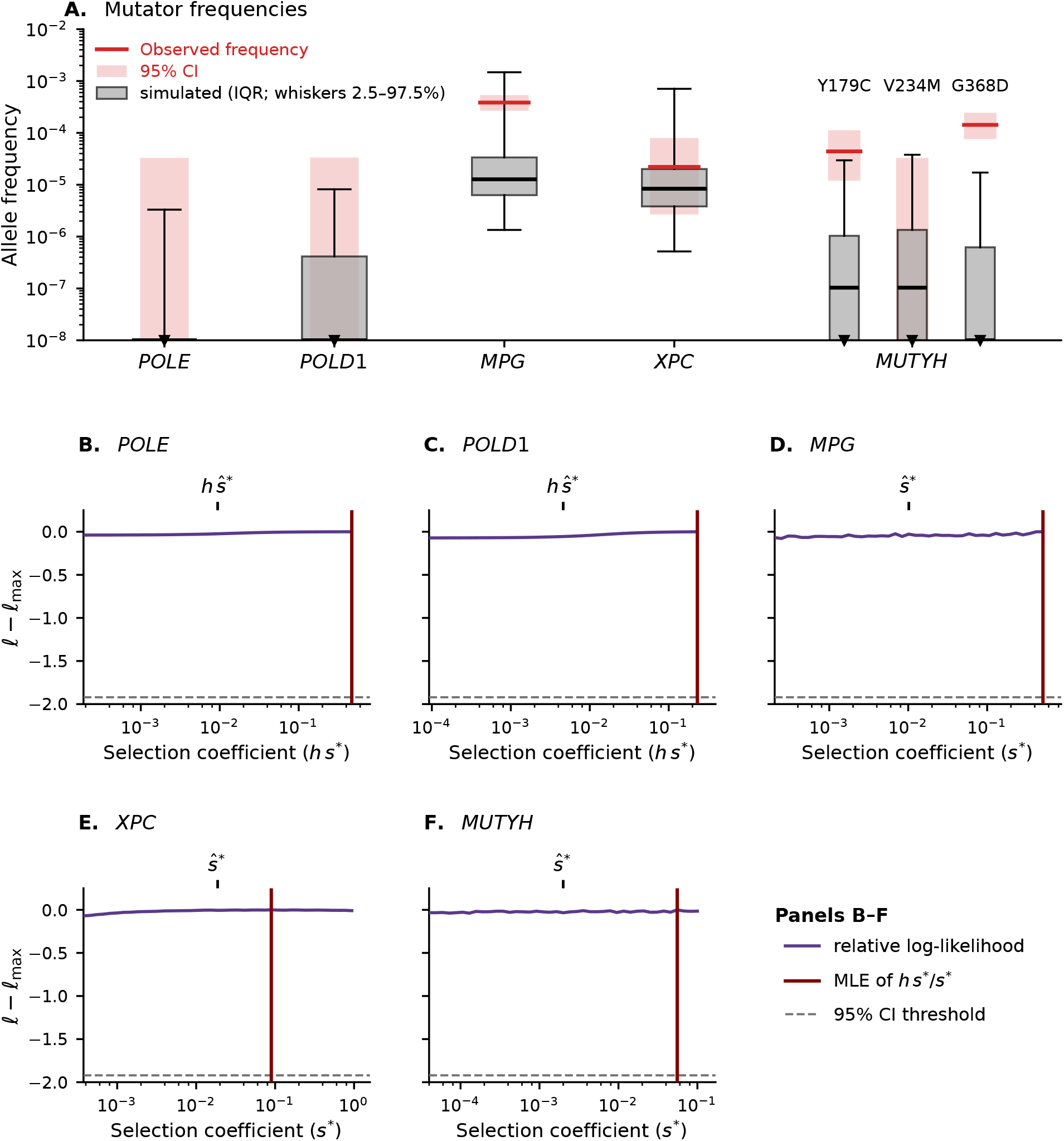
Predicted frequency distributions and likelihoods of known mutators under the SAS demographic model. **(A)** Frequency distributions at our estimate of the selection coefficient, *ŝ*\* (Table S3), shown with a box plot. **(B–F)** Relative log-likelihood of the observed allele counts,*l* − *l*_max_, as a function of the homozygous selection coefficient for the recessive mutators *MPG, XPC* and *MUTYH* or the heterozygous selection coefficient for the semi-dominant mutators *POLE* and *POLD1* (Supplementary section S5); panel **F** shows the joint likelihood of the three *MUTYH* mutators (Eq. S28). At the top of panels **B**-**F**, we show the selection coefficient in units of *ŝ*\*. Frequency distributions are obtained from single-site simulations (supplementary section S3.3) for *POLE, POLD1* and *MPG* and compound-heterozygote simulations (Supplementary section S3.4) for *XPC* and *MUTYH* with between 5 ×10^5^ and 4 ×10^6^ replicates per parameter combination (Supplementary section S5).

The exception is G368D in *MUTYH*, for which simulations indicate mutators with similar properties are expected at lower frequency than observed. We obtain similar results using the frequencies and demographic histories for other populations (Figure S8).

With the exception of *MUTYH*, known mutators were each found in a single genotype, so their dominance coefficients are unknown. For mutators in *MPG* and *XPC*, we can estimate *s*\* independently from *h* using their estimated effect size in homozygotes (thus teasing apart *h* from the compound parameter *hs*\*; Fuller et al. (2019)). Therefore, in principle, we can use our model to assess which dominance coefficients, *h*, best explain the observed frequencies of mutators. We do so using simulations similar to those above, except we hold *s*\* constant and vary the dominance coefficient *h* between 0 and 1 (see Supplementary Section S5). For both mutators, the observed frequencies are consistent with fully recessive fitness effects (Figure 3). However, there is again very little information: the confidence intervals cover much of the parameter space, and semi-dominance (*h* = 0.5) cannot be excluded. The data would more sharply favor a recessive model if these mutators were under much stronger selection than assumed (see the bottom of Figure 3).

**Figure 3:**
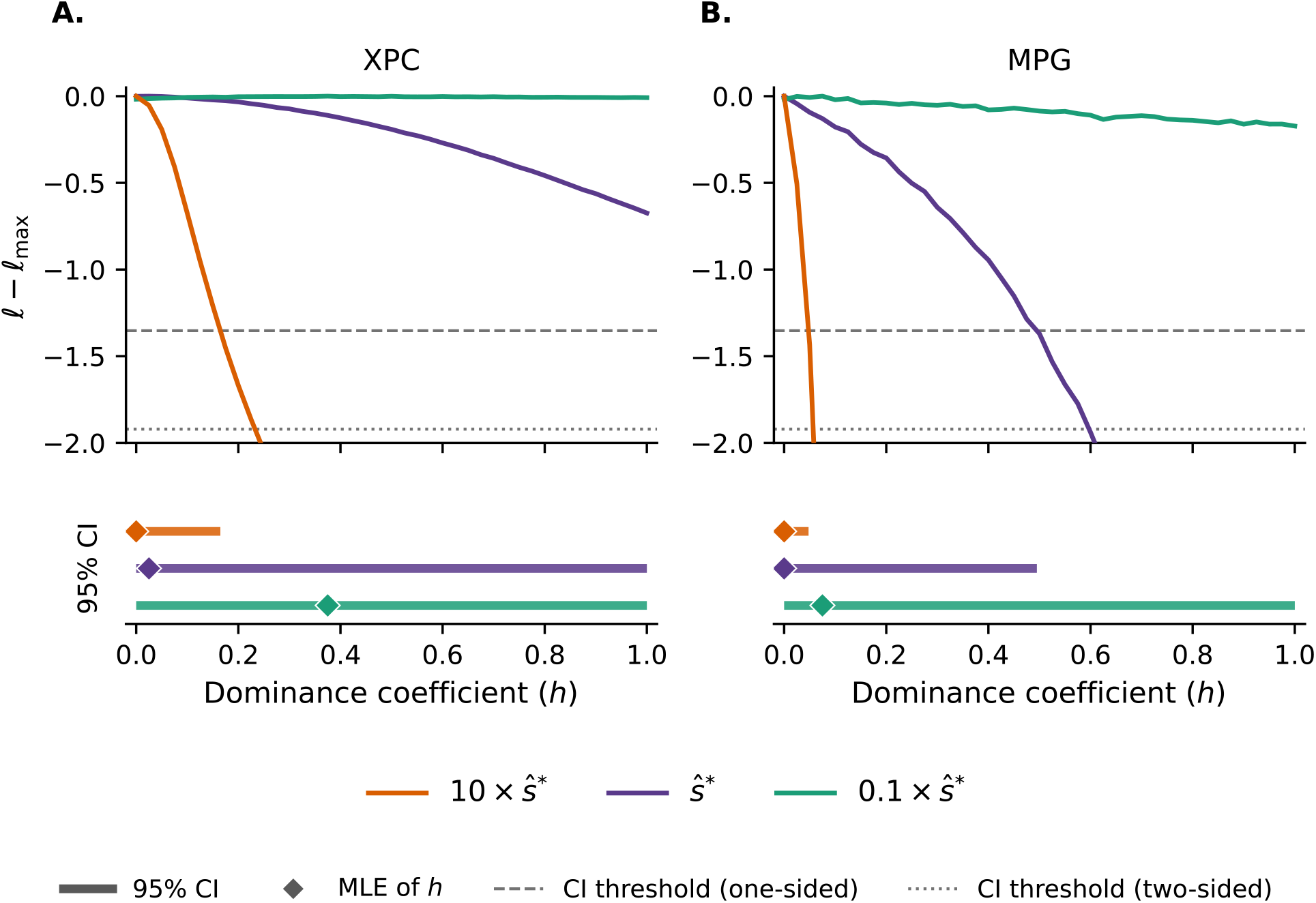
Likelihood of the dominance coefficient *h*, given the allele frequency of the mutator observed in the gnomAD v4.1.1 SAS sample. For each mutator, the selection coefficient is held constant and the relative log-likelihood of the observed allele count, *l − l*_max_, is computed across a 41-point grid of *h* ∈ [0, 1]. The log-likelihood is evaluated at *ŝ*\* (purple) (reported in Table S3), and at a ten-fold lower value 0.1 *ŝ*\* (green), and at a 10-fold higher value 10 *ŝ*\* (orange) for *XPC* (panel **A**) and *MPG* (panel **B**). Diamonds mark the maximum-likelihood estimate (MLE), and the horizontal bars below each panel denote the corresponding 95% confidence intervals. The significance threshold is 1.92 when the MLE is interior (dotted) and 1.353 when the MLE lies at the boundary *h* = 0 (dashed, see Supplementary Section S5). Likelihoods are estimated from single-site simulations for *MPG* (1 10^6^ runs per selection coefficient; Section S3.3) and compound-heterozygote simulations for *XPC* (5 10^5^ runs per selection coefficient; Section S3.4), under the SAS demographic model.

### What kind of mutators do we expect to find by surveying the number of mutations in offspring of trios?

Given that known mutators vary in their effect sizes and dominance, we investigate how these properties affect the probability of discovery. We do so by emulating the Kaplanis et al. (2022) study, in which the authors screened ~22,000 trios for offspring carrying an unexpectedly high number of DNMs, as this approach is likely to be widely followed in the future. Heterogeneity in realized DNM counts due to age and sex effects likely affects mutator discovery. To incorporate it, we assume mutators increase germline mutation rates multiplicatively and incorporate the parental age and sex effects estimated by Jónsson et al. (2017) (see Supplementary section S6). We then define a proband as an offspring who inherits twice the expected number of DNMs given their parental ages. Using this model, we ask how the probability of detecting at least one proband among a large cohort of trios, Pr(proband), depends on the genetic architecture of germline mutation rates, *i*.*e*., the number of modifier sites, their dominance coefficients, and their effect sizes.

We decompose the probability of detecting a proband into the probability that the cohort contains a mutator parent, i.e., one that expresses the mutator phenotype, and the probability that a trio with a mutator parent produces a proband. That is,

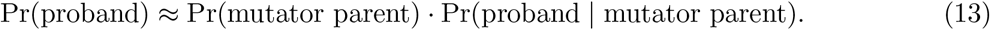

This approximation assumes that only the offspring of mutator parents could satisfy our definition of proband. Marginalized over the parental age distributions, the probability that a trio without a mutator parent produces a proband is ≈ 4.4 × 10^*−*11^ (Supplementary section S6.2), which is negligible compared to Pr(proband | mutator parent) in the range of mutator effect sizes that we consider. Below, we consider each of these terms separately.

For a mutator parent of a given sex, the probability of producing a proband behaves approximately as a step function of the mutator effect size (Figure S11). That is, below some threshold effect size, mutator parents almost never produce a proband; above it, they almost always do. The threshold corresponds to when the expected number of DNMs that an offspring inherits is doubled. Near this threshold, the probability of producing a proband gradually increases, corresponding to the Poisson variance in the realized number of DNMs. The threshold is four times smaller for fathers than mothers, because the paternal germline contributes roughly four times as many DNMs per generation as the maternal germline (Kong et al. 2012; Jónsson et al. 2017; Gao et al. 2019). For mutator effect sizes between the two thresholds, the probability of a mutator parent producing a proband without conditioning on sex reaches a plateau near 1*/*2, since the mutator is equally likely to be carried by either parent (Figure 4A). If mutator effect sizes frequently fall into this range, trio-based designs will therefore be strongly biased toward detecting mutators in fathers.

**Figure 4:**
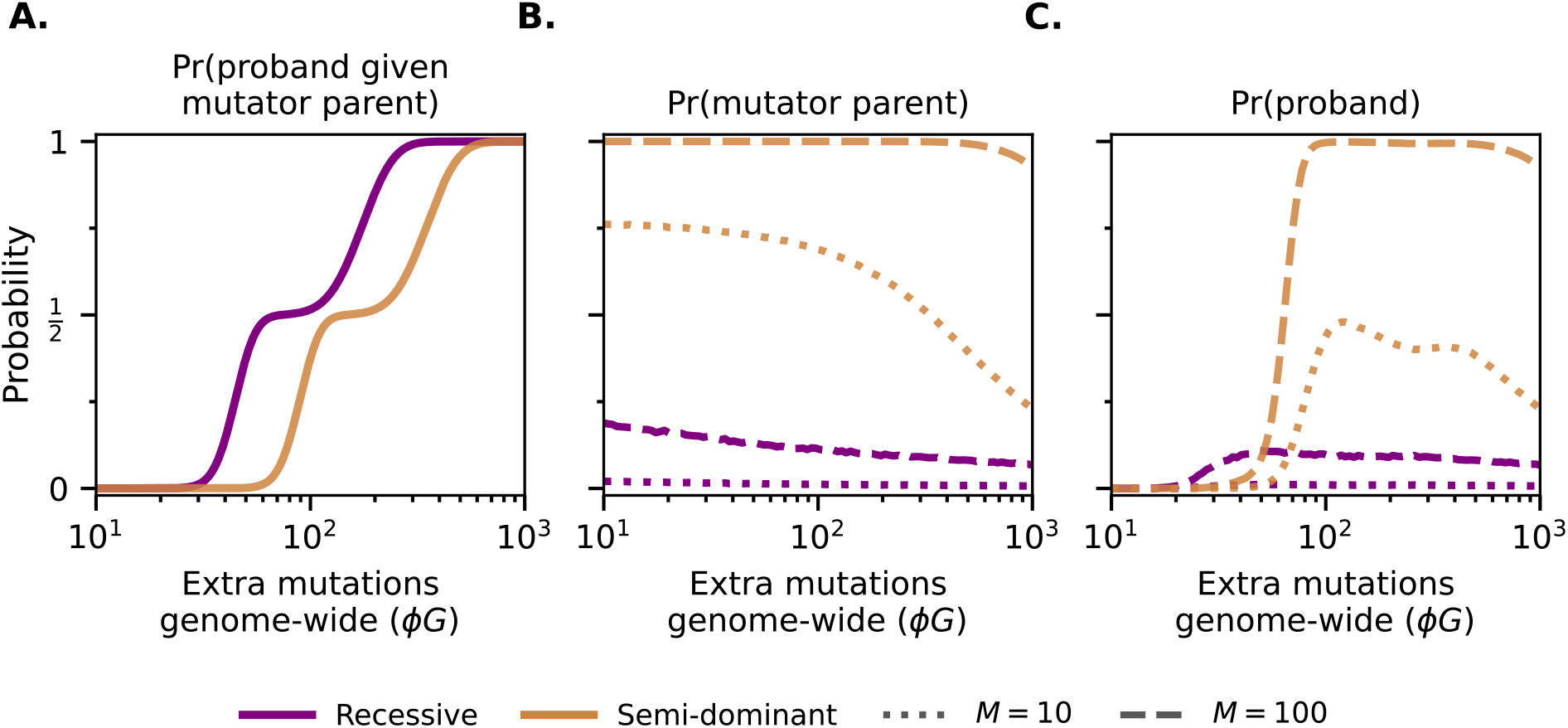
Probability of discovering a proband in a sample of ~ 22,000 trios, with sex- and age-specific parental effects on germline mutation rates following Jónsson et al. (2017) (see Supplementary section S6). **A.**Pr(proband mutator parent): for a given trio, the probability that the offspring of a mutator parent has at least twice the expected number of DNMs conditional on the parental ages. **B**. Pr(mutator parent): probability that at least one of the sampled trios includes a mutator parent. **C**. Pr(proband): probability of detecting at least one proband in the sample. Probabilities in panels B and C are estimated from 2.5 × 10^5^ replicates for each combination of effect size and dominance coefficient (see Supplementary section S6). For each parameter combination, the mutator frequency distributions are simulated using single-site simulations (see Supplementary section S3.3) under the NFE demographic model and the probabilities of a sampling a mutator parent or proband are estimated as described in Supplementary section S6. All other parameters are as described in Table 1.

The threshold effect sizes also depend on the dominance of the mutator. Mutator parents carrying partially dominant mutators are much more likely to be heterozygous, and heterozygotes express a reduced mutator phenotype compared to homozygotes. Therefore, the thresholds increase by 1*/h* for partially dominant mutators compared to fully recessive mutators, so long as selection against heterozygous mutator parents is effective (i.e., 2*N*_*e*_*hs*\* ≫ 1). Relaxing our definition of a proband shifts these thresholds down but leaves the qualitative picture unchanged, provided that parents unaffected by mutators remain unlikely to produce a proband.

The probability of detecting a proband therefore primarily reflects the probability of sampling a mutator parent carrying a mutator of sufficiently large effect. Given a mutator at a single modifier site segregating at frequency *q*, the probability that a trio contains at least one mutator parent is

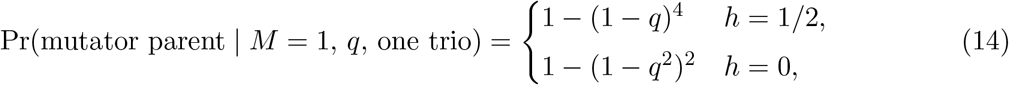

which for rare mutators is approximately 4*q* when *h* = 1*/*2 and 2*q*^2^ when *h* = 0. Therefore, for strongly selected mutators, which can only segregate at rare frequencies, the probability of sampling at least one mutator parent becomes

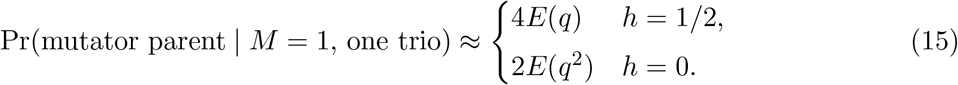

Since the frequency of mutators declines with effect size (Figure 1), so does the probability of sampling a mutator parent. For a given effect size, recessive mutators are expected to segregate at higher frequencies than partially dominant mutators. However, short of high levels of consanguinity, the probability of sampling a mutator parent is greater for partially dominant mutators, as mutator parents need only be heterozygous (as opposed to homozygous for recessive mutators). That said, for a single trio and modifier site, the probability of sampling a mutator parent is low under all conditions.

Next, we consider how the probability of sampling at least one mutator parent depends on the number of trios in the cohort. In this case, there are multiple chances to sample a mutator parent, such that the probability becomes

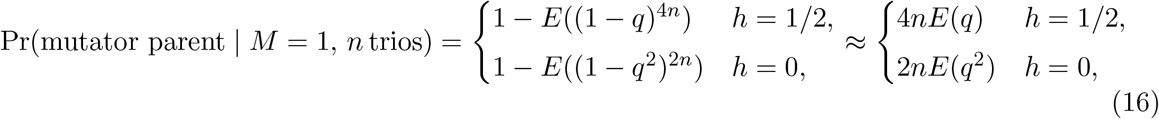

where *n* is the number of trios. These approximations assume that *q* is always much less than 1*/*4*n* for semi-dominant mutators or 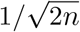 for recessive ones, which is not true for a small minority of mutators (≤ 0.1%) when *n* = 22,000. But they capture two qualitative patterns: the probability of sampling a mutator parent increases with the number of trios and remains higher for partially dominant than for recessive mutators. Nonetheless, for current study sizes, the probability of sampling a mutator parent remains low when there is only one modifier site (Figure S12).

Finally, we consider the case with many trios and multiple modifier sites. We assume that the frequencies of mutators at different modifier sites are independent and drawn from the same distribution conditional on effect size. Therefore, the probability of sampling a mutator parent becomes

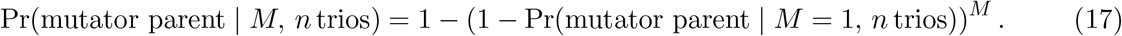

As *M* increases, so does the probability of sampling a mutator parent; for large *M*, sampling such a parent becomes nearly guaranteed.

Simulations that use a realistic demography and parental age distributions confirm these analytic results (Figure 4). In a cohort of 22,000 trios, mutator parents carrying semi-dominant mutators are much more likely to be sampled than mutator parents carrying recessive ones for a given target size. In general, many more modifier sites are needed to sample a mutator parent carrying a recessive mutator than to sample one carrying a semi-dominant mutator, regardless of effect size (Figure S6). The implication is that for effect sizes where both types of mutators can produce probands (*i*.*e*., *ϕG* ≳ 100; Figure 4A), study designs such as the one in Kaplanis et al. (2022) should be more likely to detect the offspring of mutator parents carrying semi-dominant than recessive mutators of the same effect size (Figure 4C), unless recessive mutators have a far larger target size than semi-dominant ones.

## Discussion

We study the evolution of mutators with arbitrary dominance coefficients in a panmictic population, assuming that they are under purifying selection owing to their effects in the germline. We then compare the model predictions to what is observed for seven known mutators in five genes. While the frequencies of six of the seven known mutators (all but G368D in *MUTYH*) are consistent with our model predictions, a wide range of alternative parameter values could also explain their frequencies. Similarly, for two of the modifier genes, *MPG* and *XPC*, mutator frequencies are consistent with a recessive model of selection, but we cannot exclude partial dominance. With regard to *XPC*, we note that while the mode of inheritance for the clinical phenotype associated with *XPC* dysfunction is recessive (DiGiovanna & Kraemer 2012), a previous study of LoF and other pathogenic mutations in *XPC* suggested that such mutations have a nonzero fitness effect in heterozygotes (Judd et al. 2026).

For one of the mutators, G368D in *MUTYH*, our model provides a relatively poor fit for any fitness and dominance effects (Figure 2). The same is true if we lower the average fitness cost of new deleterious mutations (Figure S7), or if we use the observed mutator frequency and demographic history estimated for Africans and African-Americans instead (Figure S8). That the model provides a poor fit in multiple ancestry groups indicates that the discrepancy is not due to ascertainment bias in a particular ancestry. Since a higher mutation rate would improve the fit, one possibility is that the mutation rate at this site is underestimated by Roulette (Seplyarskiy et al. 2023).

More broadly, our analyses of mutators are relatively uninformative about the strength or mode of selection acting against them, because they are based on the observed frequency of only one variant (for *MUTYH*, three). For mutators generated by LoF mutations, such as in *XPC*, additional information could be gleaned by jointly analyzing the frequencies of many LoF mutations at once, as they are expected to have equivalent effects on mutation rates. Similar analyses may be possible for missense mutations, accounting for potential variation in their phenotypic effects. For example, saturation mutagenesis of *MUTYH* suggests that 41% of possible missense variants impair the function of the gene (Hemker et al. 2025). If a substantial fraction of these variants elevates germline mutation rates *in vivo*, estimates of their selection and dominance coefficients could be substantially improved by considering them together. Moreover, following the approach taken here, it will be informative to compare the estimated selection coefficients to those predicted by models of selection on germline mutations alone. Such comparisons would benefit from improved estimates of the average fitness cost of new mutations (see Kar et al. (2026)).

In this regard, we note that our model and related ones (e.g., Kimura (1967), Lynch et al. (2016), and Milligan et al. (2022)) only account for the fitness costs of mutators that arise from the additional germline mutations that they cause (but see Lynch (2010)). Moving forward, drawing on models that link mutation rates to DNA replication, repair, and cell division (e.g., Spisak et al. (2024)) could help to make predictions about the effects of mutators in the soma as well. For example, mutators that act predominantly during whole genome replication may have much larger effects in dividing cell types (e.g., in spermatogonial stem cells and colonic crypts) than non-dividing ones (e.g., in oocytes and neurons). In contrast, mutators that reduce the accuracy of DNA repair may have similar multiplicative effects across cell types. Such considerations may suggest that mutators such as those in *POLE*, which increase mutation rates to a similar extent in mothers and fathers (Table S3), have effects independent of cell division. The predictions of such models could be tested empirically by observing how mutation rates in various cell types increase with age in carriers of mutators.

A further complication in modeling effects of mutators on the soma is how to translate an increase in mutation rates into a fitness cost (Zhu et al. 2025; De Manuel et al. 2026). Known germline mutators often increase somatic mutation rates, contributing to increased risk of cancers and other disorders (Robinson et al. 2021; Robinson et al. 2022). This effect likely increases their fitness cost beyond what is considered in standard models, including ours, and results in a further decrease in their frequency, their expected effect on the mean germline mutation rate in the population, and the probability that they are sampled in pedigree studies. While the qualitative effect is clear, quantification will require an estimate of the fitness effects of somatic mutations.

Even without the inclusion of such pleiotropic effects, our analyses suggest that known mutators have sufficiently large effect sizes for them to be strongly selected against. However, the broader genetic architecture of mutation rates, including the effect sizes and dominance of mutators yet to be discovered, remains poorly delimited. Our results point to a high polygenicity of mutation rates (Figure 1), which would suggest that many more subtle mutators remain to be discovered. Two straightforward ways to learn more about this architecture are to increase trio cohort sizes so that rare mutators of large effect are more likely to be sampled and to analyze pedigrees with multiple offspring in order to more precisely estimate parental mutation rates (e.g., Sasani et al. (2019)). It may also be helpful to focus on parents of a specific sex or age (Figure 4). Under the assumption that mutators increase the mutation rate multiplicatively, those carried in fathers are more likely to be detected—and indeed, the two mutators found in Kaplanis et al. (2022) were carried by fathers. Analogously, given the effects of parental ages on mutation rates, mutators carried by older parents may be easier to detect. Combining these two considerations, cohorts of older fathers may provide greatest insight into the genetic architecture of mutation rates. Additionally, methods that identify mutators through changes in the mutational spectra and not just the overall mutation rate (e.g., Young et al. (2024) and Getseva et al. (2026)) may expand the range of effect sizes that can be detected.

Even with the limited current data, however, some inferences about this architecture are already possible. The first two mutators identified by surveying trios (in *XPC* and *MPG*) seem to be recessive (Kaplanis et al. 2022). Yet their effect sizes fall in the range (*ϕG >* 100) in which we expect semi-dominant mutators to be more frequently identified. The implication is that modifier sites that produce large effect, recessive mutators outnumber those that produce large effect, semi-dominant ones by at least an order of magnitude (Figure S6). This conclusion is consistent with a broader trend across deleterious mutations, in which those of larger effect tend to be more recessive (Kacser & Burns 1981; Phadnis & Fry 2005; Agrawal & Whitlock 2011; Huber et al. 2018; Kyriazis & Lohmueller 2024). We expect further such insights into the genetic architecture of germline mutation rates to be gained by interpreting empirical discoveries of mutators in light of mathematical models of mutators.

## Data availability

This paper relied on publicly available data from the gnomAD v4.1 release. The code to run simulations, analyze data, and generate figures are available at https://github.com/matin-saeidi/mutators.

## Acknowledgments

We thank Vanessa Getseva, Hannah Munby, Lin Poyraz and other members of the Andolfatto, Przeworski and Sella labs, as well as Josh Schraiber for many helpful discussions. This work was supported by GM115889 to GS, GM153355 to MP, and F32GM163397 to WM.

