## Supplementary Materials for "Dynamics of mutators of arbitrary dominance in humans"

**S7 Additional figures** **22**

**References** **26**

### S1 Choice of notation and definitions

Table S1: Notation used throughout.

| Symbol | Meaning |
|---|---|
| *Model parameters* | |
| $N_e$ | Effective population size |
| $G$ | Haploid genome length |
| $f$ | Fraction of the genome under strong selection |
| $s_{het}$ | Heterozygous selection coefficient of a deleterious allele |
| $u_0$ | Mutation rate of individuals not carrying any mutators |
| $\mu$ | Mutation rate at a modifier site |
| $\phi$ | Mutator effect size, i.e., the increase in $u$ in homozygotes |
| $h$ | Dominance coefficient of the mutator |
| $M$ | Number of modifier sites |
| *Additional notation* | |
| $\phi G$ | Expected increase in the number of DNMs transmitted from homozygous mutator parents |
| $q$ | Mutator frequency |
| $g$ | Genotype at a modifier site (expressed as $0, 1, 2$ copies of the mutator) |
| $u$ | Germline mutation rate of an individual |
| $\hat{u}$ | Estimated mutation rate in humans |
| $\lambda$ | Expected number of additional DNMs linked to a mutator per generation |
| $s$ | Selection coefficient against mutators |
| $s^*$ | Selection coefficient against homozygous mutators when they are treated as typical deleterious alleles with a constant fitness effect |
| $\hat{s}^*$ | Our estimate of $s^*$ for a given mutator |
| $\Psi$ | Stationary distribution of mutator frequency, $\Psi(q)$ |
| $n$ | Number of trios in the cohort |
| *Notation used in the Supplement* | |
| $\Phi$ | Mutator effect size as a fold increase in mutation rate |
| $D$ | Number of DNMs inherited by an offspring |
| $Y$ | Number of DNMs passed on by a parent |
| $A_F,\ A_M$ | Age of the father and of the mother at conception |
| $\pi_F,\ \pi_M$ | Densities of the distributions of the age of fathers and mothers at conception |

### S2 Model

#### S2.1 Derivation of selection coefficient against mutators

People affected by mutators are not selected against *per se*; rather, their progeny are less fit. To model this scenario, Milligan et al. (2022) approximated the strength of selection on a mutator by calculating the expected total number of progeny descended from the person in which the mutator first arose, $K_T$. The rationale is that defining the selection coefficient, $s$, as $s = 1/K_T$ will generate the correct frequency of the mutator at mutation–selection balance (Kimura 1967). In brief, Milligan et al. (2022) showed that this approximation for $s$ can be written as

$$s \approx 1/K_T \approx s_{het} \cdot 2f\lambda, \tag{S1}$$

where $\lambda$ is the expected number of additional alleles that become linked to a mutator each generation, $f$ is the proportion of new mutations that are deleterious, and the factor of 2 reflects the expected number of generations that a mutator remains linked to a deleterious allele under free recombination. The key assumptions for this derivation are that (1) on average, mutators are not nearly lethal, such that the expected number of excess alleles linked to them reaches its steady state value and (2) the number of DNMs that a gamete receives is Poisson-distributed. We consider violations of this first assumption below. The second assumption is convenient but not strictly true. In our model, DNM counts are overdispersed relative to a Poisson distribution because the mutation rate varies across parents, though this overdispersion has little effect on our results (Figures 1 and S1A). In reality, DNM counts are also overdispersed due to non-independence between mutations (Sasani et al. 2019) and age and sex effects (Ségurel et al. 2014), neither of which we model here.

The value of $\lambda$ reflects the difference in the number of DNMs received by a gamete carrying a mutator (hereafter, mutator gamete) versus a gamete carrying an anti-mutator (hereafter, anti-mutator gamete). We revise the derivation for $\lambda$ from Milligan et al. (2022) to allow for an arbitrary dominance coefficient. Below, we consider the case of a single modifier site with a mutator segregating at frequency $q$. The derivation remains the same when there are multiple modifier sites, so long as mutators evolve independently.

The type of gamete that a parent produces and the number of DNMs inherited by a gamete depend only on the genotype of the parent, that is the number of mutators it carries, $g \in \{0, 1, 2\}$. The probability that a parent with genotype $g$ produces a mutator gamete is

$$\Pr(\text{mutator gamete} \mid g, P) = \frac{g}{2}, \tag{S2}$$

where $P$ denotes the event that a person is chosen as a parent. In turn, the probability that they produce an anti-mutator gamete is $1 - \Pr(\text{mutator gamete}|g, P) = 1 - g/2$. The number of DNMs that the gamete inherits is assumed to be Poisson-distributed with mean $Gu$, where $G$ is the genome length and $u$ is the mutation rate of the parent given their genotype, $g$, as described by Eq. 1.

The number of DNMs inherited by a mutator gamete thus depends on the probability that the gamete was produced by a homozygote or heterozygote parent. By Bayes' theorem, the probability that a parent has genotype $g$, given that it produced a mutator gamete, is

$$\Pr(g \mid \text{mutator gamete, } P) = \frac{\Pr(g)\Pr(\text{mutator gamete, } P \mid g)}{\Pr(\text{mutator gamete, } P)}. \tag{S3}$$

Assuming random mating, the proportion of the population with genotype $g$, $\Pr(g)$, follows from Hardy-Weinberg equilibrium. For the remaining terms, we note that

$$\Pr(\text{mutator gamete, } P \mid g) = \Pr(\text{mutator gamete} \mid g,\ P)\Pr(P \mid g) = \frac{g}{2}\Pr(P \mid g). \tag{S4}$$

We assume differences in mean fitness conditional on genotype are small (i.e., $s \ll 1$); as a result, the probability that a person with genotype $g$ is chosen a parent, $\Pr(P \mid g)$, is well approximated by the probability under neutrality, $1/N_e$. Therefore, Eq. S4 becomes

$$\Pr(\text{mutator gamete, } P \mid g) \approx \frac{g}{2}\frac{1}{N_e}. \tag{S5}$$

In turn, we calculate $\Pr(\text{mutator gamete, } P)$ as

$$\Pr(\text{mutator gamete, } P) = \sum_{g=0}^{2} \Pr(\text{mutator gamete, } P \mid g) \Pr(g) \approx \frac{q}{N_e}, \tag{S6}$$

where $q$ is the mutator frequency. We then approximate Eq. S3 as

$$\Pr(g \mid \text{mutator gamete, } P) \approx \frac{\Pr(g)}{q}\frac{g}{2}. \tag{S7}$$

The expected number of DNMs that a mutator gamete inherits follows from the expected number conditional on a given parental genotype, $g$, weighted by the probability that an individual has that genotype given that it produced a mutator gamete, $\Pr(g \mid \text{mutator gamete, } P)$, summed across genotypes. That is,

$$E(\text{DNMs} \mid \text{mutator gamete}) \approx \sum_{g=0}^{2} Gu \cdot \frac{\Pr(g)}{q}\frac{g}{2} = G\left(u_0 + \phi\left(q + h(1-q)\right)\right). \tag{S8}$$

Similar considerations show that

$$E(\text{DNMs} \mid \text{anti-mutator gamete}) \approx G\left(u_0 + \phi h q\right), \tag{S9}$$

which again assumes $s \ll 1$.

Given the expected number of DNMs that each type of gamete inherits, we can now derive the selection coefficient against the mutator using Eq. S1. The expected number of excess DNMs that become linked to a mutator each generation, $\lambda$, is the difference between Eq. S8 and Eq. S9, such that

$$\lambda = \phi G\big(q(1-2h) + h\big). \tag{S10}$$

Following Eq. S1, the resulting selection coefficient against mutators is

$$s \approx s_{het} \cdot 2f\phi G(q(1-2h) + h), \tag{S11}$$

which reflects the expected fitness decrease from the steady state number of excess deleterious alleles linked to the mutator.

Equation S11 reveals that the strength of selection against the mutator depends on its frequency. For our approximation of $s$ not to rely on the frequency trajectory of the mutator, we employ a separation of timescales argument, considering that the number of excess deleterious alleles linked to the mutator equilibrates much faster than its frequency changes. In Figure S1A, we show that the expected number of deleterious alleles linked to the mutator is indeed well described by Eq. S10 based on its current frequency for a range of dominance coefficients. The simulations in Figure S1A reflect our worst case scenario, where mutators initially segregate at frequency 1/2 but are not yet linked to any deleterious alleles. Otherwise, these simulations follow the general dynamics described in Supplementary section S3.1. The number of mutations linked to a mutator rapidly equilibrates and then closely tracks the equilibrium value as the frequency of the mutator changes. We also show that the average difference in fitness between individuals carrying the mutator and those who do not, $s_{\text{obs}}$, agrees with the expected selection coefficient (Eq. S11) for a range of mutator frequencies and dominance coefficients (Figure S1B).

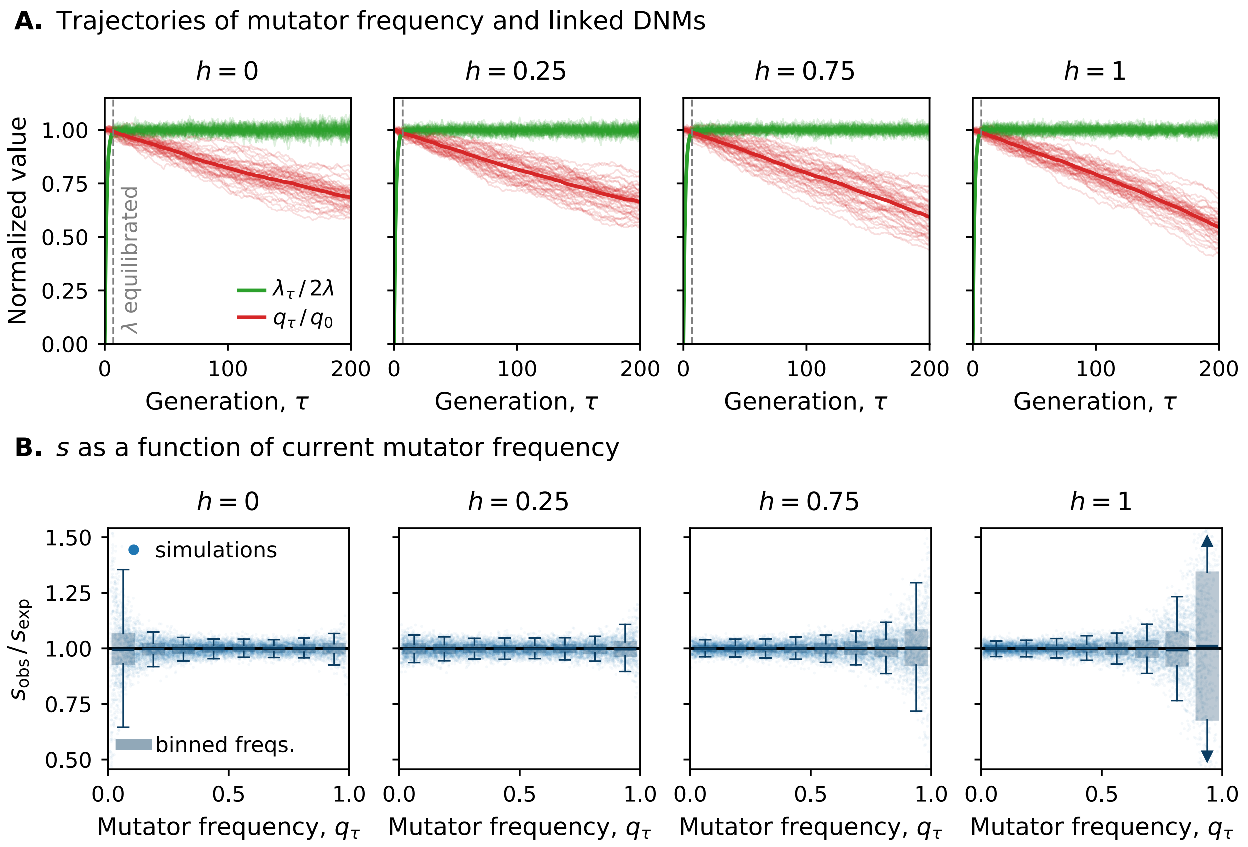

Figure S1: Validation of the selection coefficient $s$. **A.** Trajectories of mutator frequency (red) and the number of alleles linked to the mutator (green) over 200 generations. We show the mutator frequency relative to its starting value, $q_0 = 0.5$, and the number of linked alleles relative to its steady state expectation, $2\lambda$ (Eq. S10), which updates as the frequency of the mutator changes. Trajectories from 50 individual replicates are shown as thin lines, while the mean results are shown as bold lines. The dashed grey line indicates the first time that the number of alleles linked to the mutator is within 1% of its equilibrium value **B.** The average difference in fitness between individuals carrying the mutator and those who do not, $s_{\text{obs}}$, relative to the expectation given the current frequency of the mutator, $s_{\text{exp}}$ (Eq. S11), for $10^4$ replicates per dominance coefficient. Each point represents the frequency of the mutator and the fitness of individuals carrying the mutator from a single replicate simulated for 50 generations. In each panel, we bin replicates into eight equal-width bins and show the box plot per bin. All simulations use $N = 20{,}000$, $\phi G = 100$, $f = 0.08$, $u_0 = 0$, and $s_{\text{het}} = 5 \times 10^{-4}$.

#### S2.2 Selection against strongly selected mutators

The approximation of $s$ given by Eq. S1 relies on a linear approximation of the fitness function, assuming that $s \approx 2fs_{het}\lambda \ll 1$. When mutators cause a sufficiently large increase in the mutation rate, such that $\lambda \gtrsim 1/2fs_{het}$, this approximation is no longer valid. We therefore approximate the fitness of persons that carry mutators using an exponential form:

$$1 - s \approx (1 - s_{het})^{2f\lambda} \approx \exp(-s_{het} \cdot 2f\lambda). \qquad \text{(S12)}$$

This expression reduces to Eq. S11 when $s_{het} \cdot 2f\lambda \ll 1$ but accurately captures the selection against nearly lethal mutators.

##### S2.3 Derivation of the stationary distribution of mutator frequencies

We derive the stationary distribution of mutator frequencies using the diffusion approximation for a biallelic site based on the first two moments of allele frequency change (Ewens 2004). For an allele with frequency $q$, the stationary density is

$$\Psi(q) = \frac{C}{V(\Delta q)} \exp\left\{2 \int \frac{E(\Delta q)}{V(\Delta q)}\, dq\right\}, \tag{S13}$$

where $C$ is a normalizing constant. Substituting Eqs. 4 and 6 for $E(\Delta q)$ and $V(\Delta q)$ respectively, we obtain

$$\Psi(q) \approx C \exp\left\{-2N_e s^* \left[q^2 + 2hq(1-q)\right]\right\} \{q(1-q)\}^{4N_e\mu - 1}, \tag{S14}$$

after evaluating the integral and absorbing constants into $C$.

The approximation for the stationary distribution diverges at the boundaries 0 and 1 in the weak mutation regime, $4N_e\mu < 1$. To calculate the density at these states, we follow the procedure described in Ewens (2004) and used by Milligan et al. (2022). Specifically, we return to the discrete Markov chain from which the diffusion was derived and calculate the density at a given boundary state given the probability of transitioning to and from that state. The Markov chain has $2N_e + 1$ possible states, representing the possible number of mutator copies, $i$, in a finite population of size $N_e$. We approximate the density at internal states, $\Omega_i$, by integrating Eq. S14 over the interval of width $1/2N_e$ centered at $i/2N_e$,

$$\Omega_i = \int_{\frac{i-1/2}{2N_e}}^{\frac{i+1/2}{2N_e}} \Psi(q)\, dq, \qquad 1 \le i \le 2N_e - 1. \tag{S15}$$

The probability of transitioning from state $0 < i < 2N_e$ to state 0, $p_{i\to 0}$, in a single generation follows from the expected change in mutator frequency, Eq. 4 and binomial sampling, such that

$$p_{i\to 0} = \left(1 - \left(\frac{i}{2N_e} + E(\Delta q)\right)\right)^{2N_e}. \tag{S16}$$

Similar considerations show that

$$p_{i\to 2N_e} = \left(\frac{i}{2N_e} + E(\Delta q)\right)^{2N_e}. \tag{S17}$$

We assume that the probability of transitioning from one boundary state to the other is negligible, i.e., that $p_{2N_e\to 0} = p_{0\to 2N_e} \approx 0$, which is valid so long as the population size is not tiny. The boundary states can only be left via mutation, so

$$p_{2N_e\to 2N_e} = p_{0\to 0} = (1-\mu)^{2N_e}. \tag{S18}$$

At steady state, the probability flow into and out of each boundary state must balance, which allows us to calculate the density at each boundary state as

$$\Omega_0 = \frac{\sum_{i=1}^{2N_e-1} \Omega_i\, p_{i\to 0}}{1 - p_{0\to 0}}, \qquad \Omega_{2N_e} = \frac{\sum_{i=1}^{2N_e-1} \Omega_i\, p_{i\to 2N_e}}{1 - p_{2N_e\to 2N_e}}. \tag{S19}$$

The normalizing constant $C$ is chosen so that

$$\Omega_0 + \Omega_{2N_e} + \sum_{i=1}^{2N_e-1} \Omega_i = 1. \tag{S20}$$

We approximate an integral over the stationary distribution for some function of the mutator frequency, $a(q)$, as

$$\int_0^1 a(q)\Psi(q)dq \approx a(0)\Omega_0 + a(1)\Omega_{2N_e} + \sum_{i=1}^{2N_e-1} a\left(\frac{i}{2N_e}\right)\Omega_i. \tag{S21}$$

#### S2.4 Sensitivity to the choice of $s_{het}$

Our model assumes that deleterious mutations at selected sites have a fixed fitness cost, $s_{het}$. In practice, deleterious mutations vary in their fitness costs. Milligan et al. (2022) demonstrated that the dynamics of mutators are insensitive to variation in $s_{het}$ across selected sites and that selection against mutators is primarily determined by the expected fitness cost of deleterious mutations. We rely on their results and interpret $s_{het}$ as the expected selection coefficient of deleterious mutations at selected sites. However, the distribution of fitness effects for new mutations is still an open question, and uncertainty in the value of $s_{het}$ remains. To estimate reasonable values of $s_{het}$, we rely on estimates that ∼8% of the human genome is constrained, i.e., has evolved under strong purifying selection such that $2N_e s_{het} \gg 1$ (Davydov et al. 2010; Ponting & Hardison 2011; Ward & Kellis 2012; Rands et al. 2014). Given an ancestral effective population size for humans of $N_e = 2 \cdot 10^4$ (Schiffels & Durbin 2014), $s_{het}$ at these sites must be far greater than $2.5 \cdot 10^{-5}$. For our main results, we assume $s_{het} = 5 \cdot 10^{-4}$. Alternative values of $s_{het}$ do not change our qualitative results, as the same three selection regimes still appear (Fig. S2). However, if $s_{het}$ is smaller than we assume, mutators require larger effect sizes to evolve under strong selection and can generate larger increases in the mean mutation rate.

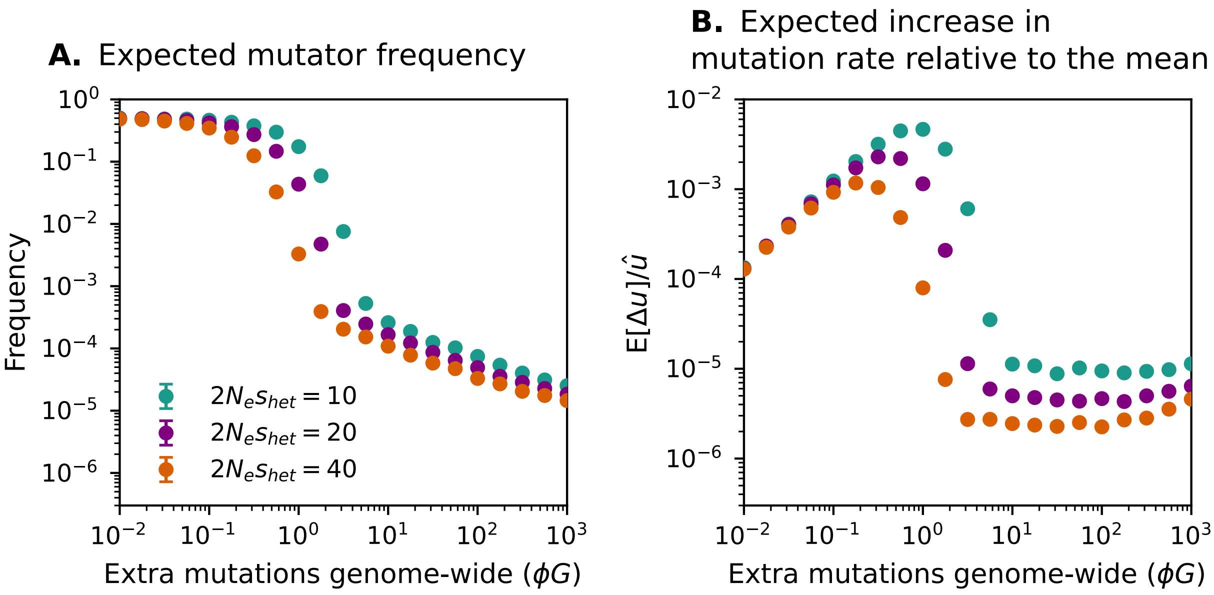

Figure S2: Sensitivity of mutator simulations to the assumed strength of purifying selection against deleterious alleles. Results are shown for recessive mutators under three different values of $2N_e s_{het}$. Otherwise, simulations are run as described in Figure 1. **A.** Expected mutator frequency as a function of mutator effect size, $\phi G$. **B.** Expected increase in the mean mutation rate relative to the estimated human point mutation rate, $\mathrm{E}[\Delta u]/\hat{u}$, as a function of $\phi G$.

### S3 Simulations

For the results presented here, we use three types of simulations. The first type, "mutator" simulations, is an adaptation of the one introduced in Milligan et al. (2022) and realizes our model in full. The second type, "single-site" simulations, is adapted from Simons et al. (2014) and simplifies our model by assuming that mutators evolve under a constant selection and dominance coefficient, as is typically assumed for deleterious alleles. The third type, "compound-heterozygote" simulations, modifies the single-site simulations and allows mutators at different modifier sites in the same gene to form compound heterozygotes that phenocopy homozygotes.

#### S3.1 Mutator simulations

Mutator simulations explicitly track the number of deleterious alleles linked to each mutator within individuals. For computational efficiency, we track only the number of deleterious alleles that an individual carries instead of their genotype at every selected site, assuming deleterious alleles are never homozygous. We initialize the genotypes at each modifier site by first sampling a mutator frequency from its stationary distribution (Eq. 7) and then assigning genotypes via binomial sampling. The initial number of deleterious alleles carried by each person is drawn from a Poisson distribution with mean $2u_0 fG/s_{het}$, which is the expectation in the absence of mutators.

We first run simulations for a burn-in period of $10N_e$ generations to allow the population to reach mutation–selection–drift balance, then continue them for $200N_e$ more generations. After burn-in, mutator frequencies are recorded every $8N_e$ generations. These recorded frequencies are then pooled across replicate simulations to obtain the mean and variance of mutator frequencies.

#### S3.2 Justification for single site simulations

Explicitly tracking the transient linkage between mutators and deleterious alleles causes mutator simulations to be prohibitively slow when the population size is large. We therefore rely on simulations that approximate mutators as deleterious alleles with a constant selection coefficient. Given a single modifier site, a selection coefficient equal to $s^* = 2Gf\phi s_{het}$ exactly reproduces the expected change in frequency in Eq. 4 under Wright-Fisher sampling with mutation and selection. The agreement between our analytic approximations that rely on Eq. 4 and results from mutator simulations (Figure 1; see also Figure 1 in Milligan et al. (2022)) suggests fluctuations in the strength of selection experienced by mutators have little impact on their expected frequency. We therefore expect the same level of agreement between the mutator and these simplified simulations. We confirm this agreement directly for recessive mutators: both types of simulation produce nearly identical moments of mutator frequency across a range of effect sizes (Fig. S3).

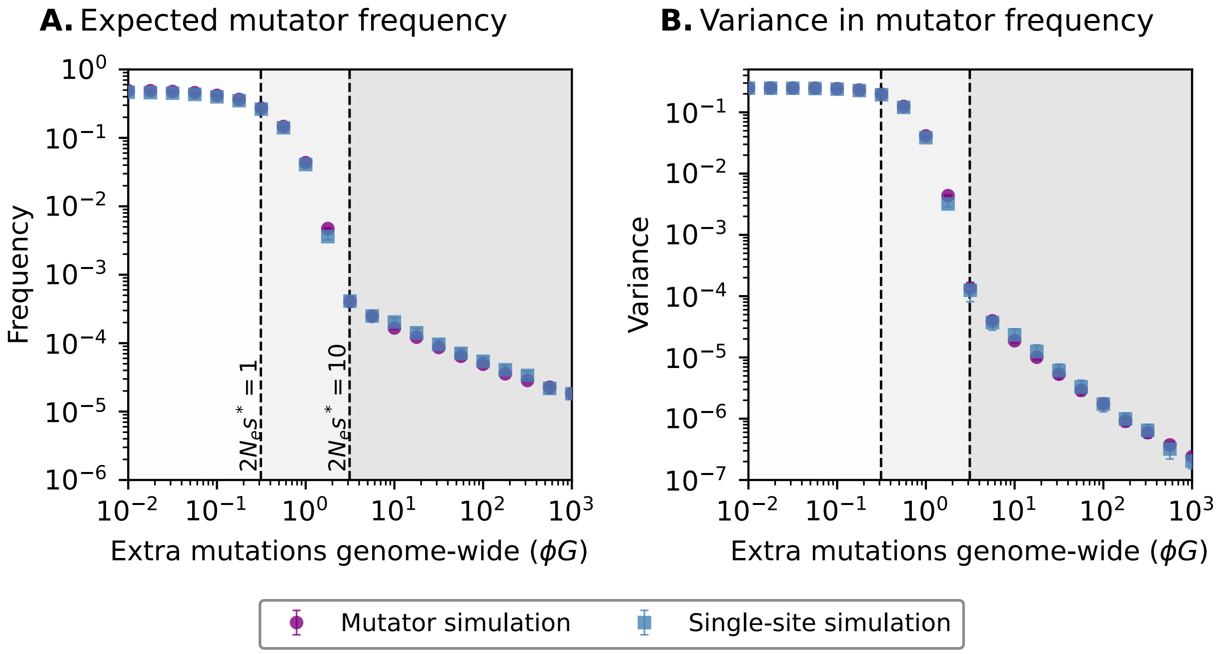

Figure S3: Comparison of mutator simulations and single-site simulations. Expected mutator frequency (**A**) and variance (**B**) as a function of the expected increase in DNMs produced by homozygous carriers of the mutator, $\phi G$. Purple circles denote results from 50 replicates of the mutator simulations with one modifier site (Supplementary Section S3.1); blue squares denote results from 100,000 replicates of the single-site simulation (Supplementary Section S3.3). Error bars indicate $\pm 2$ standard errors, though most are too small to see and therefore not shown. Both simulations assume $2N_e\mu = 5 \times 10^{-4}$ and $2fGs_{het} = 2.4 \times 10^5$, with a fully recessive mutator ($h = 0$). Dashed black lines indicate selection regime boundaries, as in Figure 1.

#### S3.3 Single-site simulations

The first type of simplified simulation, 'single-site', assumes mutators only arise from mutations at a single modifier site. At the start of the simulation, the modifier site is fixed for the anti-mutator. Each generation, new mutators can arise through forward mutation and existing mutators can revert through back mutation at rate $\mu$. We assume the number of forward and backward mutations are Poisson distributed with means $2N_e(1-q)\mu$ and $2N_e q\mu$, respectively, where $q$ is the mutator frequency. When estimating the frequency distribution of a known mutator, we use the mutation rate for the corresponding mutation type estimated in Seplyarskiy et al. (2023). Otherwise, we assume $\mu = 1.25 \cdot 10^{-8}$. After mutation, the next generation is formed by Wright–Fisher sampling with parents chosen according to their fitness. Given a realistic demographic history, we simulate the evolution of this modifier site forward in time to generate the frequency of the mutator at present. We retain every replicate, including those in which the mutator has been lost by the present and whose frequency in the population is therefore zero.

We also use these simulations to estimate the ages of known mutators (Figure S10). Because the mutator may have arisen from multiple independent mutations, we report its age based on when the oldest surviving copy arose. Therefore, we record the generation in which a forward mutation (i.e., one that creates a mutator) arises and track the number of mutator copies descended from it. In these simulations, we consider only replicates in which the mutator is still segregating in the population at present.

##### S3.4 Compound-heterozygote simulations

Mutations at multiple sites in *MUTYH* and *XPC* are either observed or strongly suspected to cause the mutator phenotype in compound heterozygotes (Clark 1998; Lehmann et al. 2011; Kaplanis et al. 2022; Sherwood et al. 2023; Young et al. 2024). In the 'compound-heterozygote' simulations, we allow for multiple modifier sites within a gene but assume that each haplotype carries at most one mutator. This assumption lets us treat the entire gene as a single biallelic locus with mutation, selection, and drift. The compound-heterozygote simulations are therefore equivalent to the single-site simulations but with a higher mutation rate. Because we neglect the possibility that a forward mutation lands on a haplotype that already carries a mutator, we may overestimate the frequency of haplotypes that carry mutators. When mutators are rare, this bias should be negligible.

These simulations are initialized and parameterized in the same manner as the single-site simulations, except for the mutation rate at modifier sites. For *MUTYH*, we assume that only the three missense variants (Y179C, V234M, and G368D) identified as mutators to date may complement each other; the forward mutation rate is taken as the sum of the mutation rates to these three variants. For *XPC*, we assume that all LoF mutations can complement each other and thus use the total LoF mutation rate across the gene (see Supplementary Section S4.2).

These simulations do not track the site at which a forward mutation occurred or the variant that it produced. In order to obtain the frequency of our focal variants, we randomly assign each forward mutation to a specific variant with a probability proportional to the mutation rate of the variant relative to the total mutation rate.

##### S3.5 Demographic models

We simulate mutator frequencies under three demographic models, corresponding to those inferred for the non-Finnish European (NFE), South Asian (SAS), and African and African-American (AFR) genetic ancestry groups (Figure S4). Each model is a composite of two sources: recent estimates capturing the explosive population growth of the last few hundred generations, and older estimates for the more distant past. For the recent past, we use the estimates obtained from Schraiber et al. (2025) for NFE or from Kar et al. (2026) for SAS and AFR. Beyond the oldest timepoint provided by each study, we use the effective population sizes inferred by Schiffels and Durbin (2014) for the corresponding population: CEU (Utah residents with Northern and Western European ancestry) for NFE, GIH (Gujarati Indians in Houston, Texas) for SAS, and YRI (Yoruba in Ibadan, Nigeria) for AFR. Schiffels and Durbin (2014) report times in years, which we convert to generations using the generation time of 30 years that they assume.

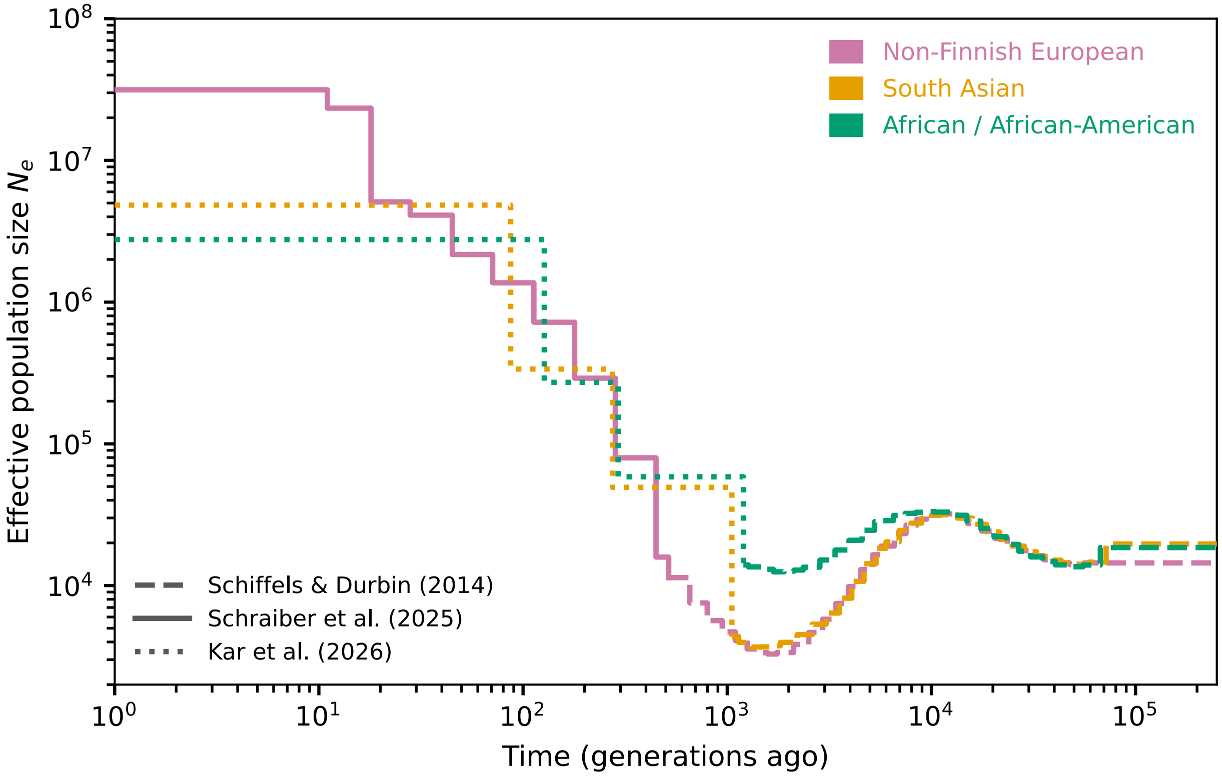

Figure S4: The demographic histories of the non-Finnish European (NFE), South Asian (SAS), and African and African-American (AFR) ancestry groups. The dotted lines represent the effective population sizes inferred by Kar et al. (2026); the solid line represents those inferred by Schraiber et al. (2025); and the dashed lines represent those inferred by Schiffels and Durbin (2014). See Supplementary section S3.5 for more details.

### S4 Known mutators in humans

Table S2: Population frequencies of known human mutators.

| Population | *XPC* p.Arg220Ter | *MPG* p.Ala135Thr | *POLE* p.Leu424Val | *POLD1* p.Ser478Asn | *MUTYH* p.Tyr179Cys | *MUTYH* p.Gly368Asp | *MUTYH* p.Val234Met |
|---|---|---|---|---|---|---|---|
| European non-Finnish | $1.36 \times 10^{-5}$ | $1.80 \times 10^{-4}$ | $8.47 \times 10^{-7}$ | 0 in 1,172,856 | $2.33 \times 10^{-3}$ | $5.88 \times 10^{-3}$ | $3.17 \times 10^{-4}$ |
| European Finnish | $4.69 \times 10^{-5}$ | 0 in 63,152 | 0 in 63,886 | 0 in 62,800 | $1.61 \times 10^{-3}$ | $1.82 \times 10^{-3}$ | 0 in 63,976 |
| East Asian | 0 in 44,854 | $6.68 \times 10^{-5}$ | 0 in 44,892 | 0 in 44,180 | 0 in 44,888 | $6.68 \times 10^{-5}$ | 0 in 44,892 |
| South Asian | $2.21 \times 10^{-5}$ | $3.84 \times 10^{-4}$ | 0 in 91,090 | 0 in 89,328 | $4.39 \times 10^{-5}$ | $1.43 \times 10^{-4}$ | 0 in 91,086 |
| Middle Eastern | 0 in 6,076 | $1.16 \times 10^{-3}$ | 0 in 6,060 | 0 in 6,036 | $1.65 \times 10^{-4}$ | $4.95 \times 10^{-4}$ | 0 in 6,084 |
| African | 0 in 74,910 | $1.33 \times 10^{-4}$ | 0 in 75,050 | 0 in 74,602 | $3.33 \times 10^{-4}$ | $9.19 \times 10^{-4}$ | $1.33 \times 10^{-5}$ |

Mutator frequencies in each genetic ancestry group, as reported in gnomAD v4.1.1. Entries of the form "0 in $n$" indicate that the mutator was not observed among the $n$ chromosomes sampled in that group. The *POLD1* mutator (rs397514632) is absent from gnomAD v4.1.1, so we take its sample sizes from the immediately adjacent 3' site, rs1199543659 (chr19:50,406,457, GRCh38).

#### S4.1 Estimating mutator effect sizes

The studies that discovered mutators in humans reported their effects on germline mutation rates in different ways; here, we place their effect sizes on a comparable quantitative scale. Our aim is to estimate the effect of mutators on the mutation rate of the parent, as distinct from the total number of DNMs inherited by the offspring. Additionally, we want effect sizes that reflect the averages across parental ages and sex.

To that end, we ask how many DNMs the mutator parent transmitted, relative to the number expected of a parent of the same sex and age with no mutators. The number transmitted, $Y$, is not observed directly, because only a subset of the DNMs of an offspring can be assigned to a parent of origin. For each trio, we instead observe the total number of DNMs in the child, $D$, and the numbers phased to the father and to the mother. Assuming that whether a DNM is phased is independent of its parent of origin, we impute the number transmitted by the mutator parent as

$$\tilde{Y} = D \cdot \frac{\text{DNMs phased to the mutator parent}}{\text{DNMs phased to either parent}}. \tag{S22}$$

We compare $\tilde{Y}$ to $E(Y \mid A_F)$ or $E(Y \mid A_M)$, the number of DNMs expected from a father or mother of that age in the absence of mutators, which we predict from the age- and sex-effects estimated in Jónsson et al. (2017), such that

$$E(Y \mid A_F) = 6.05 + 1.51\, A_F, \qquad E(Y \mid A_M) = 3.61 + 0.37\, A_M, \tag{S23}$$

where $A_F$ and $A_M$ are the ages of the father and mother at conception. The multiplicative effect size of the mutator, $\Phi$, then follows as

$$\hat{\Phi} = \frac{\tilde{Y}}{E(Y \mid A_F)} \quad \text{or} \quad \hat{\Phi} = \frac{\tilde{Y}}{E(Y \mid A_M)}, \tag{S24}$$

for fathers or mothers, respectively. We average $\hat{\Phi}$ across parents that carry the mutator, assuming that they are independent. When multiple children from the same parent were sequenced, we first average across offspring to generate a more precise estimate of $\Phi$ in that parent. For *MUTYH*, we discard the one offspring with no DNMs phased to the germline of the mutator parent, given that there is insufficient information.

Finally, we translate the multiplicative effect size into the average increase in the mutation rate, $\phi$, that the mutator causes in homozygotes. Assuming the mutator parent is homozygous, the two definitions are related as follows:

$$\phi G = \frac{1}{2}(\Phi - 1)\left[\int E(Y \mid A_F)\,\pi_F(A_F)\,dA_F + \int E(Y \mid A_M)\,\pi_M(A_M)\,dA_M\right] = \frac{1}{2}(\Phi - 1)\,E(D), \tag{S25}$$

where $G$ is the genome size and $\pi_F$ and $\pi_M$ are the densities of the distributions of paternal and maternal ages at conception (Eqs. S29 and S30). In practice, we use $E(D) = 70.3$ as the expected number of DNMs that a child inherits (Jónsson et al. 2017). If the mutator parent is heterozygous, we replace the left-hand side of Eq. S25 with $h\phi G$.

This approach makes three somewhat unrealistic assumptions. First, we neglect that the different approaches for sequencing, DNM calling, and phasing taken by these studies likely affect the total number of DNMs detected or the fraction that are phased. However, given the small number of trios for each mutator, error in our effect size estimates is likely dominated by the randomness in the DNM counts rather than by methodological differences among studies. Second, we assume that the three mutators in *MUTYH* have the same effect size, even though missense mutations may vary in their phenotypic effects. Indeed, the G368D mutator appears to have a larger effect than the V234M mutator (Table S3). We nonetheless make this assumption for simplicity and because none of the trios we analyze include a homozygote from which to estimate effect sizes independent of other mutators. Finally, we assume that mutators cause the same fold increase in maternal and paternal mutation rates, which may not be true, depending on how and when during development the mutators introduce additional mutations. Among the sequenced trios, only mutators in *MUTYH* and *POLE* were carried by both sexes. For *MUTYH*, the mutator father had a slightly smaller multiplicative increase than the mutator mothers, which is consistent with the hypothesis from (Young et al. 2024) that these mutators generate excess C>A mutations during early embryonic development. For *POLE*, both sexes had similar multiplicative increases in their mutation rates (Sherwood et al. 2023).

Table S3: Estimates of mutator effect sizes.

| Gene | Offspring ID | Genotype | Mutator parent | Age | Phased Pat./Mat. | $D$ | $E(Y)$ | $\hat{\Phi}$ | $\bar{\hat{\Phi}}$ | $\phi$ | $\phi G$ | $\hat{s}^*$ |
|---|---|---|---|---|---|---|---|---|---|---|---|---|
| *XPC* | GEL_1 | R220*/R220* | Father | 30–35 | 129 / 1 | 425 | 55.1 | 7.65 | 7.65 | $7.8 \times 10^{-8}$ | 234 | 0.019 |
| *MPG* | GEL_3 | A135T/A135T | Father | 35–40 | 87 / 5 | 306 | 62.7 | 4.62 | 4.62 | $4.2 \times 10^{-8}$ | 127 | 0.010 |
| *POLE* | A:II.1 | | | 23 | 11 / 23 | 93 | 12.1 | 5.19 | | | | |
| | A:II.2 | L424V/+ | Mother | 25 | 11 / 19 | 119 | 12.9 | 5.86 | | | | |
| | A:II.3 | | | 30 | 23 / 14 | 156 | 14.7 | 4.01 | | | | |
| | B:III.1 | L424V/+ | Mother | 22 | 15 / 14 | 109 | 11.8 | 4.48 | 4.34 | $3.9 \times 10^{-8}$ | 117 | 0.0094 |
| | B:III.2 | | | 24 | 14 / 9 | 84 | 12.5 | 2.63 | | | | |
| | B:II.1 | L424V/+ | Father | 22 | 41 / 2 | 158 | 39.3 | 3.84 | | | | |
| | B:II.2 | | | 24 | 51 / 4 | 231 | 42.3 | 5.07 | | | | |
| *POLD1* | A:II.1 | | | 19 | 11 / 7 | 56 | 10.6 | 2.05 | | | | |
| | A:II.2 | S478N/+ | Mother | 22 | 9 / 9 | 60 | 11.8 | 2.55 | | | | |
| | A:II.3 | | | 25 | 14 / 12 | 65 | 12.9 | 2.33 | 2.64 | $1.9 \times 10^{-8}$ | 58 | 0.0046 |
| | B:II.1 | S478N/+ | Mother | 19 | 15 / 12 | 76 | 10.6 | 3.17 | | | | |
| | B:II.2 | | | 24 | 6 / 6 | 69 | 12.5 | 2.76 | | | | |
| *MUTYH* | B:II.1 | Y179C/G368D | Mother | 23 | 7 / 11 | 71 | 12.1 | 3.58 | | | | |
| | B:II.2 | | | 26 | 4 / 11 | 57 | 13.2 | 3.16 | | | | |
| | C11 | Y179C/V234M | Mother | 20 | 11 / 7 | 49 | 11.0 | 1.74 | | | | |
| | C12 | | | 24 | 20 / 4 | 71 | 12.5 | 0.95 | | | | |
| | C21 | | | 34 | 9 / 2 | 49 | 16.2 | 0.78 | 1.71 | $8.3 \times 10^{-9}$ | 25 | 0.0020 |
| | C23 | Y179C/V234M | Mother | 32 | 12 / 6 | 54 | 15.4 | 1.48 | | | | |
| | C22 | | | 36 | 5 / 0 | 34 | 16.9 | –[a] | | | | |
| | C31 | Y179C/V234M | Father | 36 | 14 / 6 | 79 | 60.4 | 0.92 | | | | |
| | C32 | | | 33 | 19 / 4 | 72 | 55.9 | 1.07 | | | | |
| | C41 | V234M/+ | Mother | 32 | 14 / 1 | 46 | 15.4 | 0.41 | | | | |
| | C42 | | | 36 | 17 / 7 | 56 | 16.9 | 1.98 | | | | |

Information used to estimate the effect size for each mutator. Each row represents one offspring. Siblings share a pedigree label (e.g., A:II) and are separated from other families by light grey lines. The genotype of the mutator parent is described by the amino acid changes; + indicates the reference allele. The expected number of DNMs passed on by the mutator parent in the absence of mutators, $E(Y)$, is calculated from Eq. S23 given the sex and the age of the mutator parent at conception. For *XPC* and *MPG*, we use the midpoint of the provided age range. The resulting effect size estimates are calculated as described in Supplementary Section S4.1; $\bar{\hat{\Phi}}$ is the average effect size for each mutator; $\hat{s}^*$ is calculated from Eq. 5 with the estimated effect size, assuming the parameter values in Table 1. [a] Offspring C22 is excluded from the estimated effect size for mutators in *MUTYH*, because no DNMs were phased to the maternal germline.

Table S4: Point mutation rates to the known human mutators.

| Gene | Variant | dbSNP | Chr. | Position | Trinucleotide context | Mutation rate $\mu$ (bp$^{-1}$ gen$^{-1}$) |
|---|---|---|---|---|---|---|
| *XPC* | R220* | rs745679643 | 3 | 14,165,549 | `T[C>T]G` | $7.92 \times 10^{-8}$ |
| *MPG* | A135T | rs200449939 | 16 | 83,139 | `G[C>T]G` | $1.04 \times 10^{-7}$ |
| *POLE* | L424V | rs483352909 | 12 | 132,673,664 | `T[C>G]T` | $3.70 \times 10^{-9}$ |
| *POLD1* | S478N | rs397514632 | 19 | 50,406,456 | `G[C>T]T` | $7.05 \times 10^{-9}$ |
| *MUTYH* | Y179C | rs34612342 | 1 | 45,332,803 | `G[T>C]A` | $1.00 \times 10^{-8}$ |
| | G368D | rs36053993 | 1 | 45,331,556 | `A[C>T]C` | $7.66 \times 10^{-9}$ |
| | V234M | rs200165598 | 1 | 45,332,479 | `A[C>T]A` | $1.13 \times 10^{-8}$ |

Genomic coordinates are given in GRCh38; dbSNP gives the reference SNP identifier of the variant. In the trinucleotide context, the mutated base is written as a pyrimidine (C or T), flanked by its 5′ and 3′ neighbors.

#### S4.2 Effect of compound heterozygosity on *XPC* variant frequency

We estimate the rate of LoF mutations in *XPC* that arise from both single nucleotide mutations and indels (Table S5). For single nucleotide mutations, we sum the Roulette mutation rate estimates (Seplyarskiy et al. 2023) across sites annotated by the Variant Effect Predictor (VEP) (McLaren et al. 2016) as stop-gained, start-lost, stop-lost, splice-donor, or splice-acceptor variants. The contribution from indels is the length of the canonical coding sequence (2823 bp) times the estimated mutation rate to indels that cause frameshifts, $8.25 \times 10^{-10}$ per site per generation (Porubsky et al. 2025).

Table S5: The LoF mutation rate of the *XPC* gene.

| Consequence | Opportunities | Mutation rate (gen$^{-1}$) |
|---|---|---|
| Stop-gained (nonsense) | 371 | $1.47 \times 10^{-6}$ |
| Start-lost | 9 | $4.15 \times 10^{-8}$ |
| Stop-lost | 8 | $1.20 \times 10^{-8}$ |
| Splice-donor | 90 | $2.92 \times 10^{-7}$ |
| Splice-acceptor | 90 | $2.89 \times 10^{-7}$ |
| Subtotal | 568 | $2.11 \times 10^{-6}$ |
| Frameshifts | 2,823 | $2.33 \times 10^{-6}$ |
| Total LoF | — | $4.44 \times 10^{-6}$ |

Varying the LoF mutation rate has little effect on the frequency distribution of the focal *XPC* mutator, when we use the realistic demography for the SAS population (Figure S5A). This insensitivity is expected given that the recent explosion in effective population size causes most mutators to be young and therefore to have experienced little selection either as a homozygote or as a compound heterozygote. In contrast, under a constant population size of $N_e = 10^6$, increasing the LoF mutation rate lowers the frequency distribution of the focal mutator (Figure S5B).

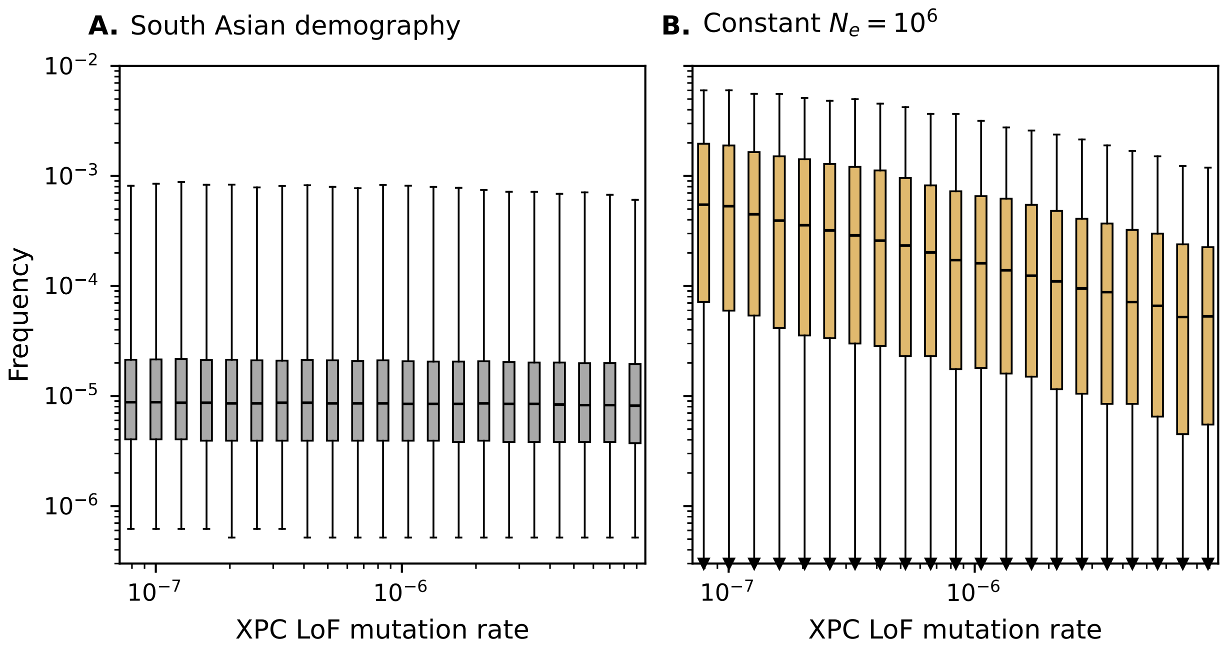

Figure S5: Frequency distributions of the *XPC* mutator for different rates of LoF mutations. In panel **A** are results for the SAS demographic model (Section S3.5) from 100,000 replicates; in **B** for a constant population-size model with $N_e = 10^6$ from 5,000 replicates (due to computational intensity). Otherwise, simulations are run as in Figure 2A.

### S5 Likelihood estimation and tests of neutrality

#### S5.1 Likelihood estimates for $s^*$

We estimate the likelihood of $s^*$ via Monte Carlo estimation for each mutator. At each tested value of $s^*$, we simulate the frequency of the mutator at present under the SAS demographic history (Supplementary section S3.5), using single-site simulations (Supplementary section S3.3) or, for the mutators in *XPC* and *MUTYH*, compound-heterozygote simulations (Supplementary section S3.4). In each case we run $R$ replicates, with $R$ between $5 \times 10^5$ and $4 \times 10^6$ according to the mutation rate of the mutator, so that a comparable number of replicates retain it across mutators. The likelihood of a given value of $s^*$, $\mathcal{L}(s^*)$, is the probability of observing the $k$ mutator copies found among the $\eta$ sequenced chromosomes in the South Asian ancestry group within gnomAD v4.1.1. We estimate this likelihood by averaging across replicates, such that

$$\mathcal{L}(s^*) = P(k \mid \eta, s^*) = \frac{1}{R} \sum_{i=1}^{R} \binom{\eta}{k} q_i^k \, (1 - q_i)^{\eta - k}, \tag{S26}$$

where $q_i \in [0, 1]$ is the mutator frequency in replicate $i$. We then construct an approximate 95% confidence interval using the likelihood-ratio statistic,

$$\mathrm{LR}(s^*) = 2\left[\, \ell_{\max} - \ell(s^*) \,\right], \qquad \ell(s^*) = \log \mathcal{L}(s^*), \tag{S27}$$

under the assumption that the LR is asymptotically $\chi^2$-distributed with one degree of freedom (Wilks 1938). We caution that since the copy number of the mutator represents only a single observation of the underlying population frequency, this approximation may be unreliable.

For *MUTYH*, we assume that the three known mutators share the same selection coefficient, allowing us to combine information across them to jointly estimate $s^*$. For simplicity, we further assume that the three variants evolve independently. The joint likelihood is therefore the product of the marginal likelihood, $\mathcal{L}_v(s^*)$ (Eq. S26), for each variant, $v$, such that

$$\mathcal{L}(s^*) = \prod_v \mathcal{L}_v(s^*). \tag{S28}$$

The variants are simulated jointly (Section S3.4), but assuming independence should be inconsequential, as the degree of compound heterozygosity appears to have little effect on the predicted frequency distributions (Figure S5A).

##### S5.2 Testing neutrality

We also ask whether the observed frequency of each mutator is consistent with neutrality, i.e., $s^* = 0$. Because $s^*$ is constrained to be non-negative, $s^* = 0$ lies on the boundary of the parameter space, so in this case, we assume that the likelihood-ratio statistic (Eq. S27) follows a mixture of $\chi^2$ distributions with zero or one degrees of freedom, $\frac{1}{2}\chi_0^2 + \frac{1}{2}\chi_1^2$ (Self & Liang 1987). In all cases, we cannot reject neutrality (Table S6), as expected given that the likelihoods are nearly flat across the range of selection coefficients considered (Figure 2B–F).

Table S6: Tests of neutrality for known mutators.

| Gene | LR | $p$ |
|---|---|---|
| *POLE* | 0.08 | 0.39 |
| *POLD1* | 0.15 | 0.35 |
| *MPG* | 0.21 | 0.33 |
| *XPC* | 1.19 | 0.14 |
| *MUTYH* | 0.07 | 0.40 |

##### S5.3 Likelihood estimates for $h$

We estimate the likelihood for the dominance coefficient, $h$, in the same manner as we do for $s^*$. When the maximum likelihood estimate is $h$=0, we proceed as described in the Section S5.2 to obtain an approximate confidence interval.

#### S6 Simulations of proband discovery in trios

We simulate the discovery of probands among a large cohort of trios, while varying the genetic architecture of mutation rates, i.e., the number of modifier sites, their effect sizes, and their dominance coefficients. These simulations consist of two stages. The first generates the frequency distribution of mutators at present using the single-site simulations (Supplementary section S3.3) under the NFE demographic history (Supplementary section S3.5). For a given combination of mutator effect size and dominance coefficient, we run 2,000,000 single-site simulations and record the frequency of each mutator at present.

During the second stage, we perform 250,000 replicates where we construct parental genotypes and estimate the probability of observing a proband. In each replicate, we sample $M$ mutator frequencies, $q_1, ..., q_M$ from the simulated frequency distribution. Parental genotypes are assigned independently at each modifier site by binomial sampling, $g_{i,j} \sim \text{Binomial}(2, q_j)$. Given the genotype of a parent, we can calculate their mutation rate. If we neglect age and sex effects, their mutation rate follows from Eq. 1, where we adjust $u_0$ so that the realized mean mutation rate equals $\hat{u}$. The number of DNMs that the parent passes down is then Poisson distributed with mean $Gu$.

However, age and sex effects can affect the probability of observing a proband. To account for them, we model mutators as multiplicatively increasing the number of DNMs passed down by a parent. In the absence of mutators, we rely on the estimates from Jónsson et al. (2017) to calculate the expected number of DNMs passed down by a parent of a given sex and age (see Eq. S23). We assume that paternal and maternal ages, $A_F$ and $A_M$ respectively, are drawn from truncated normal distributions using the mean, variance, and range of ages observed in Jónsson et al. (2017), such that

$$A_F \sim \text{TruncatedNormal}(\text{mean} = 32.0,\ \sigma = 9.0;\ \text{min} = 17,\ \text{max} = 71), \tag{S29}$$

$$A_M \sim \text{TruncatedNormal}(\text{mean} = 28.2,\ \sigma = 6.5;\ \text{min} = 16,\ \text{max} = 47), \tag{S30}$$

We convert the additive effect size of a mutator into a multiplicative effect size, $\Phi$, by rearranging Eq. S25, such that

$$\Phi = 1 + \frac{2\phi G}{E(D)}, \tag{S31}$$

and use $E(D) = 70.3$ for the expected number of DNMs that a child inherits (Jónsson et al. 2017). Therefore, each modifier site where a parent is homozygous increases the expected number of DNMs that they pass on by a factor $\Phi$ and those where they are heterozygous increase it by a factor of $1 + h\frac{2\phi G}{E(D)}$. We normalize the mean number of DNMs that a child inherits using a multiplicative correction such that mean averaged across parental ages remains constant.

Finally, we calculate the probability that a child meets our definition of our proband. Under our assumptions, the number of DNMs inherited by offspring, $D$, is Poisson-distributed with mean equal to the sum of the expected count from each parent given their age and sex. We define a proband as a child that inherits at least twice the expected number of DNMs given their parental ages, as we consider that their parents are good candidates for mutator carriers. The threshold value, $D_{\text{thr}}$, will vary across trios. Instead of explicitly sampling DNM counts for each offspring, we evaluate the probability that a child inherits more than $D_{\text{thr}}$ DNMs analytically

$$\rho = \Pr(D \geq D_{\text{thr}}) \text{ where } D \sim \text{Pois}(E(D \mid A_F, A_M)). \tag{S32}$$

The probability that at least one proband is present in the cohort is then calculated as

$$\Pr(\text{at least one proband}) = 1 - \prod_{i=1}^{n}(1 - \rho_i), \tag{S33}$$

where $i$ indexes trios and $n = 22,000$ is the number of trios.

##### S6.1 Number of modifier sites required to sample a mutator parent

The number of modifier sites required to sample a mutator parent in a cohort of $n$ trios depends on the effect size and dominance of the mutator. Given $M$ modifier sites, we estimate the probability of sampling a mutator parent as

$$Pr(\text{mutator parent}) \approx 1 - (1 - \Pr(\text{mutator parent} \mid M = 1,\ n \text{ trios}))^M, \tag{S34}$$

where $Pr(\text{mutator parent} \mid M = 1, n \text{ trios})$ is calculated using Eq. 16 and the simulations that underlie Figure 4A. In Figure S6, we show the number of modifier sites required to sample a mutator parent with probability 0.95. Across the range of effect sizes that we consider, sampling a mutator parent requires a much larger number of modifier sites when mutators are recessive than when they are semi-dominant. This pattern holds regardless of whether the number of trios is on par with the number analyzed in Kaplanis et al. (2022) or ten times that amount.

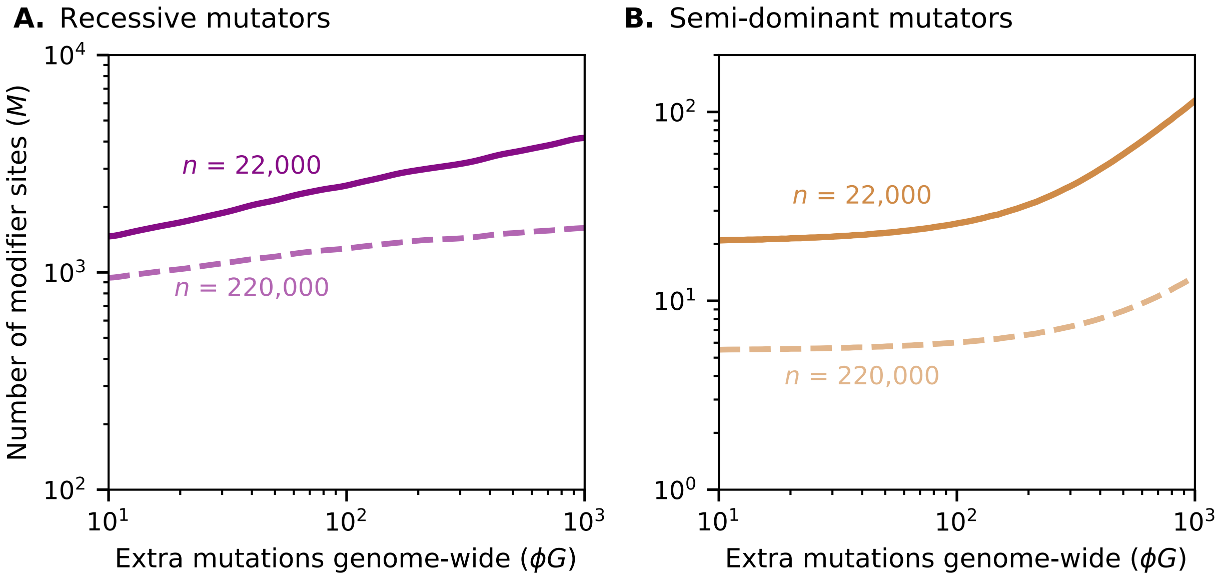

Figure S6: Number of modifier sites required for a cohort to contain at least one mutator parent with probability 0.95, as a function of the mutator effect size $\phi G$, for recessive (**A**) and semi-dominant (**B**) mutators. Solid curves, $n = 22{,}000$ trios; dashed curves, $n = 220{,}000$. $\Pr(\text{mutator parent} \mid M = 1)$ (Eq. 16) is evaluated over the simulated distribution of the mutator frequency under the NFE demographic model.

#### S6.2 Probability of producing a proband in the absence of mutators

The decomposition in Eq. 13 neglects the possibility of a proband born to parents with no mutators. Here, we show that such probands are in fact extremely unlikely. We assume that the numbers of DNMs transmitted by a father of age $A_F$ and a mother of age $A_M$ are Poisson distributed with means $E(Y \mid A_F)$ and $E(Y \mid A_M)$, respectively (Eq. S23). The number of DNMs that a child inherits, $D$, is then Poisson distributed around the sum of the paternal and maternal means, such that

$$D \sim \text{Pois}\left(E(Y \mid A_F) + E(Y \mid A_M)\right). \tag{S35}$$

Given our definition, the child must inherit more than $D_{\text{thr}}(A_F, A_M) = \lceil 2E(D \mid A_F, A_M) \rceil$ DNMs to be labelled a proband. By integrating over $\pi_F$ and $\pi_M$ (Eqs. S29 and S30), we find that the probability of a trio without mutator parents producing a proband is

$$\begin{aligned} \Pr(\text{proband} \mid \text{no mutator parent}) &= \iint Pr(D > D_{\text{thr}}) \cdot \pi_F(A_F)\, \pi_M(A_M)\, dA_F\, dA_M \\ &\approx 4.4 \times 10^{-11}. \end{aligned} \tag{S36}$$

Across a cohort of $n = 22{,}000$ trios, the probability that any proband arises without a mutator parent is $1 - (1 - \Pr(\text{proband} \mid \text{no mutator parent}))^n \approx 9.6 \times 10^{-7}$.

#### S7 Additional figures

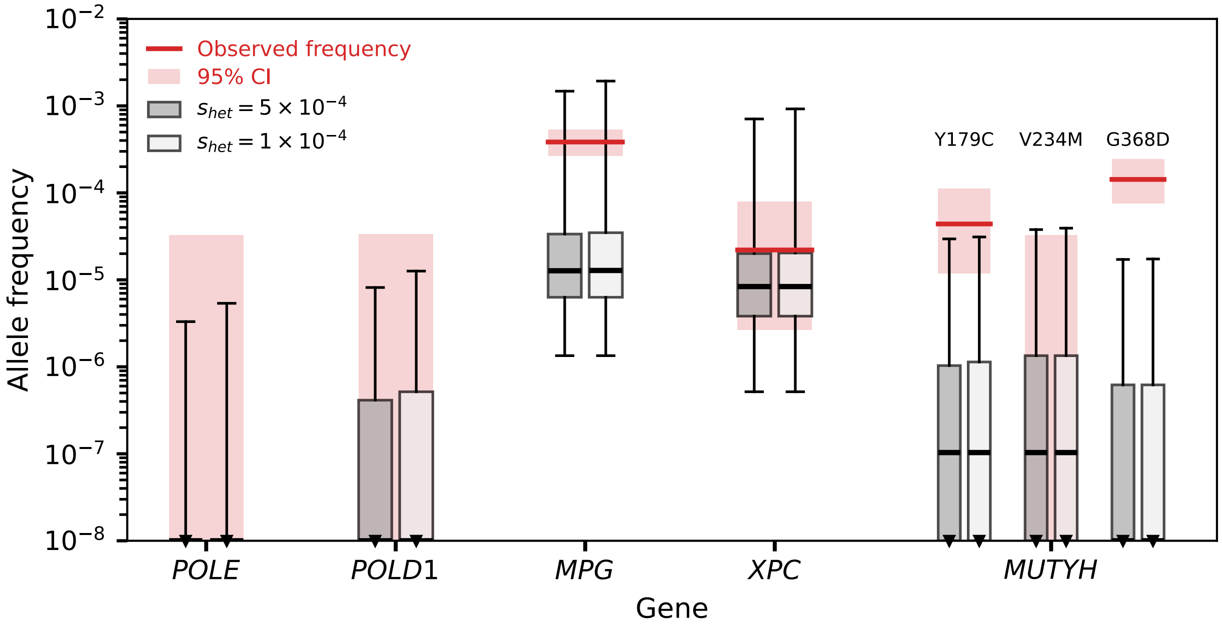

Figure S7: Frequency distributions of known mutators under our model when deleterious alleles have a five-fold lower fitness cost, $s_{het} = 1 \times 10^{-4}$. All other details are as in Figure 2A.

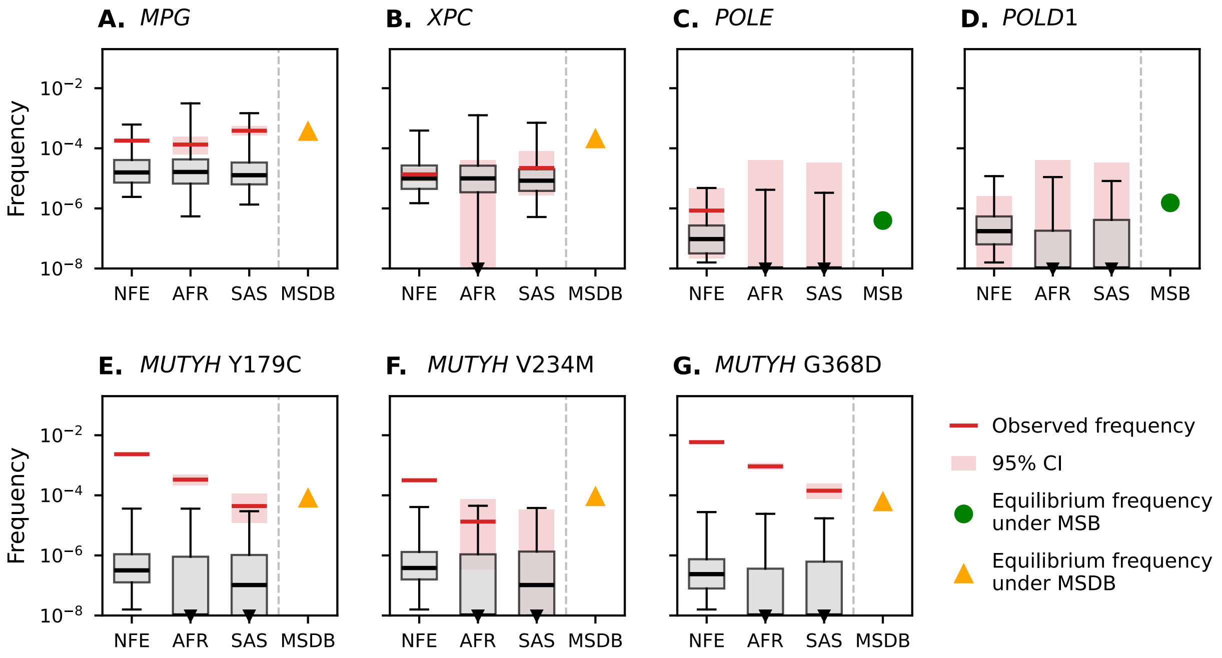

Figure S8: Predicted frequency distributions of each known mutator under the NFE, AFR, and SAS demographic models. The NFE distributions are conditional on the mutator segregating, since these mutators were identified (and therefore must be segregating) in populations of recent European ancestry. The final column shows the expected frequency under mutation–selection balance (MSB, green circle) or mutation–selection–drift balance (MSDB, orange triangle) assuming $N_e = 20{,}000$, depending on whether the mutator is assumed to be semi-dominant or recessive (Haldane 1927; Nei 1968). The simulated frequency distributions are generated in the same manner as in Figure 2A other than the choice of demographic history (see Section S3.5).

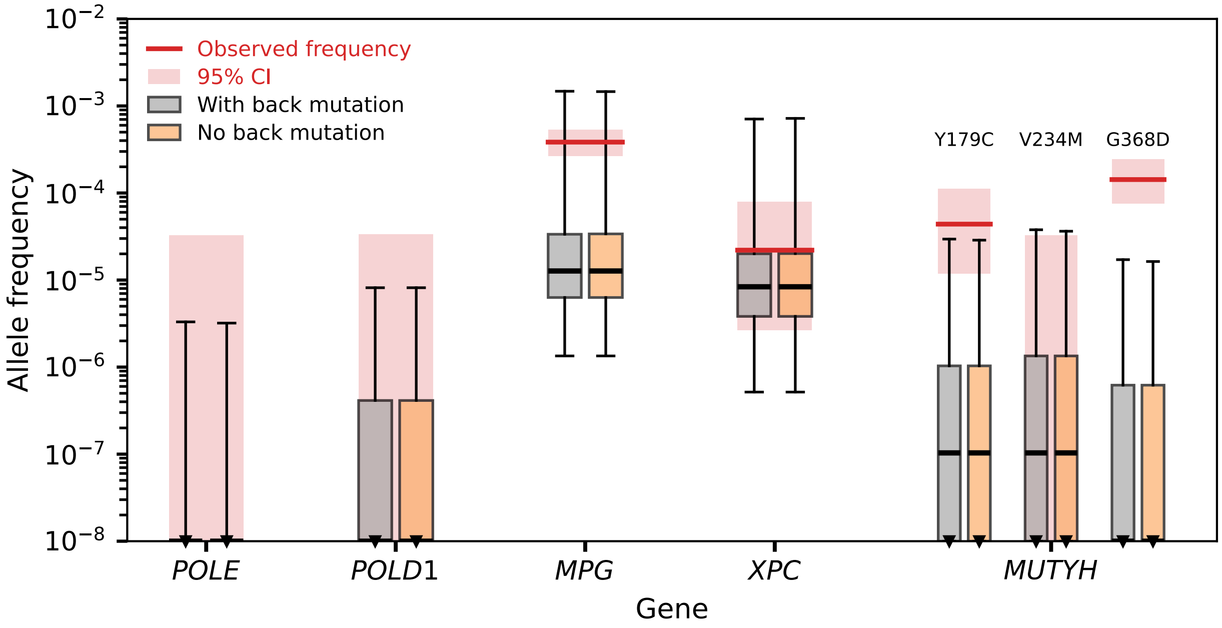

Figure S9: Predicted mutator frequency distributions with back mutation occurring at the same rate as forward mutation (grey) and with the back mutation rate set to zero (light orange). All other details are as in Figure 2A.

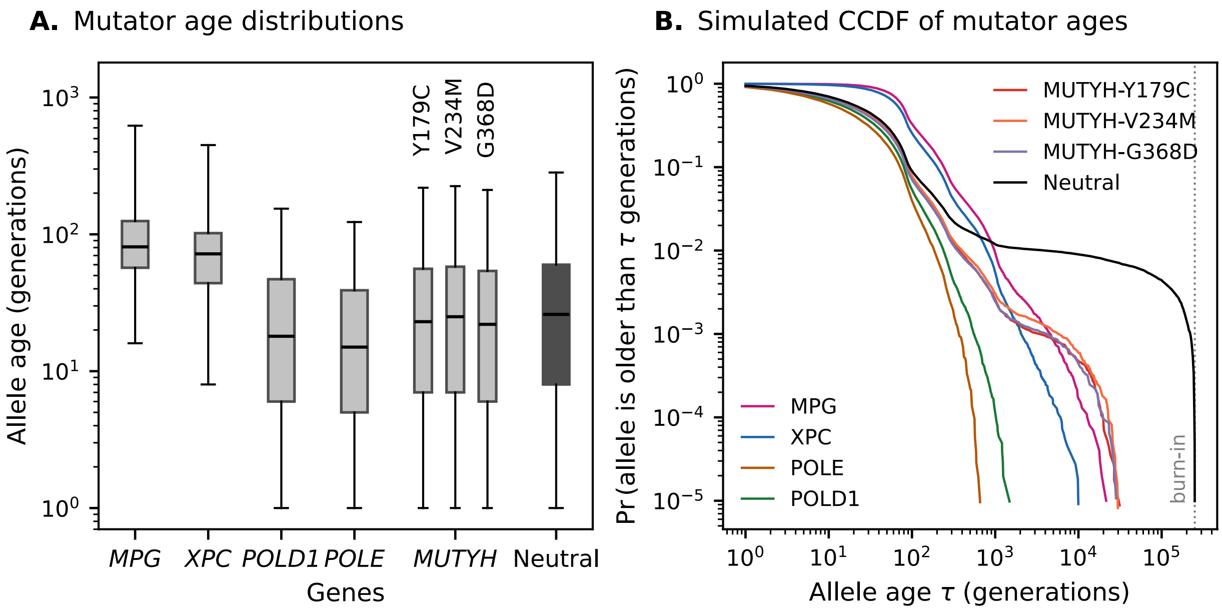

Figure S10: Age distributions of known mutators and of a neutral allele under the SAS demographic model, conditional on segregating in the present generation (Supplementary section S3.3). For the neutral allele, we assume $\mu = \hat{u}$; all other details are as in Figure 2A. **A.** Distribution of allele ages in generations, with the neutral allele in black. **B.** Complementary cumulative distributions (CCDF) of mutator ages, $\Pr(\text{Age} > x)$.

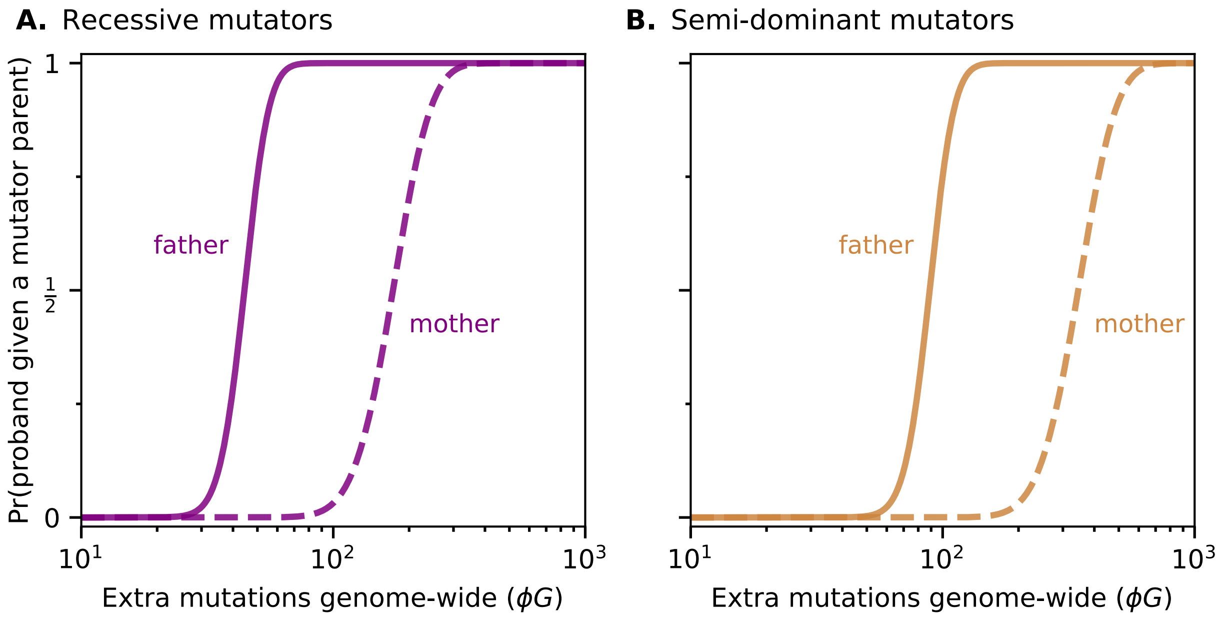

Figure S11: Probability of a proband in a trio in which exactly one parent has the mutator. All other details are as in Figure 4A.

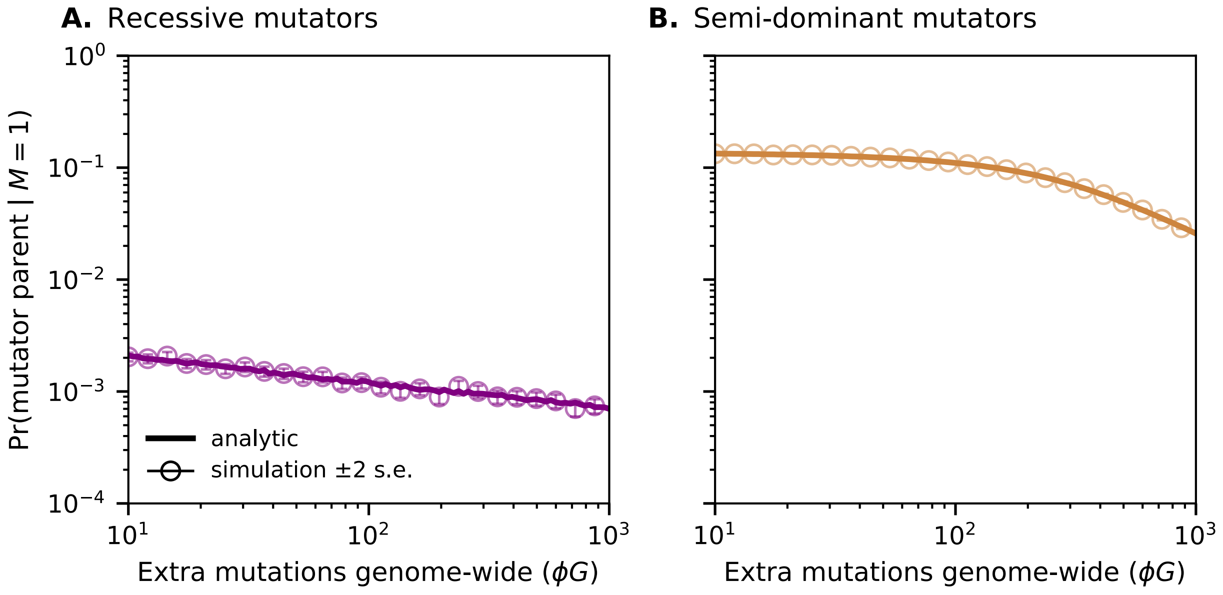

Figure S12: Probability that a cohort of $n = 22{,}000$ trios contains at least one mutator parent, given a single modifier site. The solid line shows the exact form of Eq. 16; open circles show the simulated estimates from 250,000 replicates per effect size, with error bars indicating $\pm 2$ standard errors. All other details are as in Figure 4.
